# Cross-assay RNA modeling reveals cancer biomarkers

**DOI:** 10.64898/2026.04.30.722009

**Authors:** Hope A. Townsend, Kimberly R. Jordan, Rebecca J. Wolsky, Lucy B. Van Kleunen, Natalie R. Davidson, Kian Behbakht, Matthew J. Sikora, Robin D. Dowell, Aaron Clauset, Benjamin G. Bitler

**Author notes:** **Corresponding Author:** Benjamin G. Bitler. Research Complex 2 12700 East 19^th^ Avenue Room 3400D, MS 8613 Aurora, CO 80045.

## Abstract

The clinical heterogeneity of cancer poses a major challenge for precision medicine. Limited cohort sizes across evolving assay platforms impede reliable biomarker discovery. Here, we systematically evaluate how to integrate data from four transcriptomics platforms: bulk and single-cell (sc) RNA sequencing (RNA-seq), NanoString, and microarray for predictive modeling in cancer. We use high-grade serous carcinoma (HGSC) of tube-ovarian origin as a model system, as it is highly heterogeneous in both biology and assay data.

We find that using fold-change of gene expression in patients with matched pre- and post-neoadjuvant chemotherapy samples reduces inter-patient and inter-assay variability but is insufficient to overcome platform-specific biases. Microarray and scRNA-seq data exhibit systematic biases, while RNA-seq and NanoString show the most promise for combination into a single training cohort. To mitigate inter-assay limitations, we generate a new data set of HGSC tumor samples profiled with both RNA-seq and NanoString, and use it to identify the limits of detection and optimal harmonization strategies. Our approaches enable integration of cohorts for separate and combined RNA-seq and NanoString predictive models of disease recurrence (test-set AUROCs > 0.8), validated in external microarray cohorts.

We leverage single-cell and bulk RNA-seq network-based analyses to provide mechanistic context for genes in the predictive models. Our models indicate that *GBP4* expression is a key predictor of recurrence and marks immune remodeling towards cytotoxicity. We provide an interactive web portal to facilitate exploration of data and results. These findings guide cross-assay harmonization of transcriptomic data and enable improved predictive modeling in heterogeneous cancers.

**Statement of Significance:** We present a framework for integrating RNA-seq, NanoString, microarray, and single-cell transcriptomic data for predictive modeling, enabling robust biomarker discovery in heterogeneous cancers and identifying *GBP4* as a marker of immune remodeling.

## Introduction

Precision medicine aims to assign the most promising treatment regimen based on a patient’s specific pathology (e.g. tumor microenvironment) as measured in biological data (e.g. RNA levels)[1]. Machine learning models can then be trained on these data to predict clinical endpoints, including disease recurrence, therapeutic response, and survival. Predictive models serve a dual purpose: in addition to estimating treatment success, the genes selected by the model can highlight potential biological mechanisms underlying disease progression and therapy resistance.

To build a robust predictive model, the training data must contain sufficient samples to represent the full heterogeneity of a given disease. However, that is often not the case, particularly for rare diseases or small data sets[2]. Therefore, analyses that combine smaller cohorts have often been favored to identify biomarkers less confounded by spurious correlations or batch effects that bias single-cohort studies[3,4]. While a large, integrated cohort is highly effective, it requires that all cohorts use comparable measurements (e.g., the same high-throughput sequencing assays), or a secondary transformation that can harmonize distinct measurements.

Continuous technological development has confounded this requirement by dispersing high-throughput transcriptomics data across assays that measure RNA in distinct ways, e.g., RNA-sequencing (single-cell (sc) or bulk), microarray, and NanoString. Microarray was one of the first platforms, enabling highly specific quantification even with degraded RNA by using molecular probes that hybridize to a predetermined gene set[5]. RNA-seq has dominated recent transcriptomics studies; a sample of RNA is sequenced and mapped back to a reference genome for quantification, allowing consideration of any genes/isoforms, though often with lower sensitivity and higher sampling biases[6]. As a cheaper intermediate, NanoString is a probe-based assay that avoids technical biases from post-RNA extraction steps at the cost of considering a maximum of 1,000 genes at a time[7]. Microarrays have been used to analyze large cohorts with long-term follow-up, but have evidence of limited dynamic range due to their reliance on fluorescence intensity[8]. Single-cell (sc) RNA-seq considers cell-state granularity yet effectively captures only the most highly expressed genes[9]. To compare scRNA-seq to bulk versions, expression values are usually summed within clustered cell-types to create “pseudobulk” values. The technical diversity of these assays has prevented a multi-cohort combination of data sets, despite the potential value for more powerful predictive modeling. Previous work has shown general consistency across these assays in detecting differential expression patterns[10,11]. However, studies have not yet assessed the validity of combining cohorts of different assay types for predictive modeling, despite the potential to yield more robust models.

As a result, the question of how to retain maximum data for predictive modeling with the rapid advancement of technologies remains unanswered. We address this question by interrogating assay-based biases for a complex disease that presents both patient and assay heterogeneity. High-grade serous carcinoma of tube-ovarian origin (HGSC) is the most common and aggressive subtype of epithelial ovarian carcinoma and is the fifth leading cause of cancer-death in women[12]. It is typically diagnosed at advanced stages (III/IV), where 5-year survival rates range from 20% to 40%[12]. HGSC has a largely standard treatment, including neoadjuvant chemotherapy (NACT), where patients are treated with chemotherapeutics (e.g., carboplatin/cisplatin and paclitaxel) to ideally reduce tumor size for more optimal surgical debulking. Several studies have attempted to identify biomarkers of prognosis with NACT treatment, but with limited success and overlap of results[1,13,14]. This variability in findings is often attributed to disease heterogeneity and low sample sizes[2]. Analyzing matched pre- and post-chemotherapy (pre-post) data can help address heterogeneity, as using expression changes rather than absolute values can alleviate simple scale differences from subject- and possibly assay-specific biases. However, prior work has shown that even when using such matched pre-post data, HGSC patient heterogeneity remains high, indicating a specific need for increasing sample size by combining data sets across assays[15].

In this work, we evaluate how to use matched pre- and post-chemotherapy data across different transcriptomic assays for predictive modeling in cancer, focusing on assay limitations and strengths, both individually and combined. First, we assess the limitations and comparability of results with these assays in HGSC independently. Although using matched pre-post data improved the overlap of expression trends across assays compared to using expression alone, it is still unable to fully resolve assay-specific biases. Unlike microarray and scRNA-seq, NanoString and RNA-seq do not show systemic technical biases (e.g., low dynamic range or sensitivity). Therefore, second, we create a new data set of post-NACT tissue samples assayed with both NanoString and RNA-seq. With this data set, we predict limits of detection and introduce preprocessing steps to optimally combine the assays. Third, we successfully combine NanoString and RNA-seq for predictive modeling of HGSC recurrence, identifying biomarkers that are also verifiable in both scRNA-seq and microarray data. Finally, we use the newer assays RNA-seq and scRNA-seq to predict the mechanistic context of identified biomarkers with network analyses. Overall, this work provides critical insights and data on harmonizing data between transcriptomic assays for predictive modeling, with a focus on HGSC.

## Materials and Methods

Additional and more detailed methods are in Supplemental Methods.

## Cohorts

### Longitudinal, single-assay pre-post cohort

We hypothesized that although exact expression levels differ between assays, the change in expression – measured by log2 fold-change (log2FC) – between two samples using the same assay would be consistent across assay types, since assay-specific biases would be accounted for by the ratios. Therefore, we collected transcriptomics data for paired pre- and post-NACT samples assayed with microarray, NanoString, or RNA-seq (both bulk and single-cell) with previously published clinical data (Figure 1A). Summary statistics for the combined and mini-cohorts are in Table 1. We found one pre-post cohort for microarray (N=28)[16], and three each for RNA-seq and NanoString (sample size ranging from 15 to 35) [1,13,15,17-23]. All NanoString cohorts used the same PanCancer IO 360 gene panel. Zhang et al[14] published paired pre- and post-NACT scRNA-seq and platinum-free interval (PFI) for 11 patients, 5 of whom were matched with bulk RNA-seq in an above cohort[22,23].

**Figure. 1.**
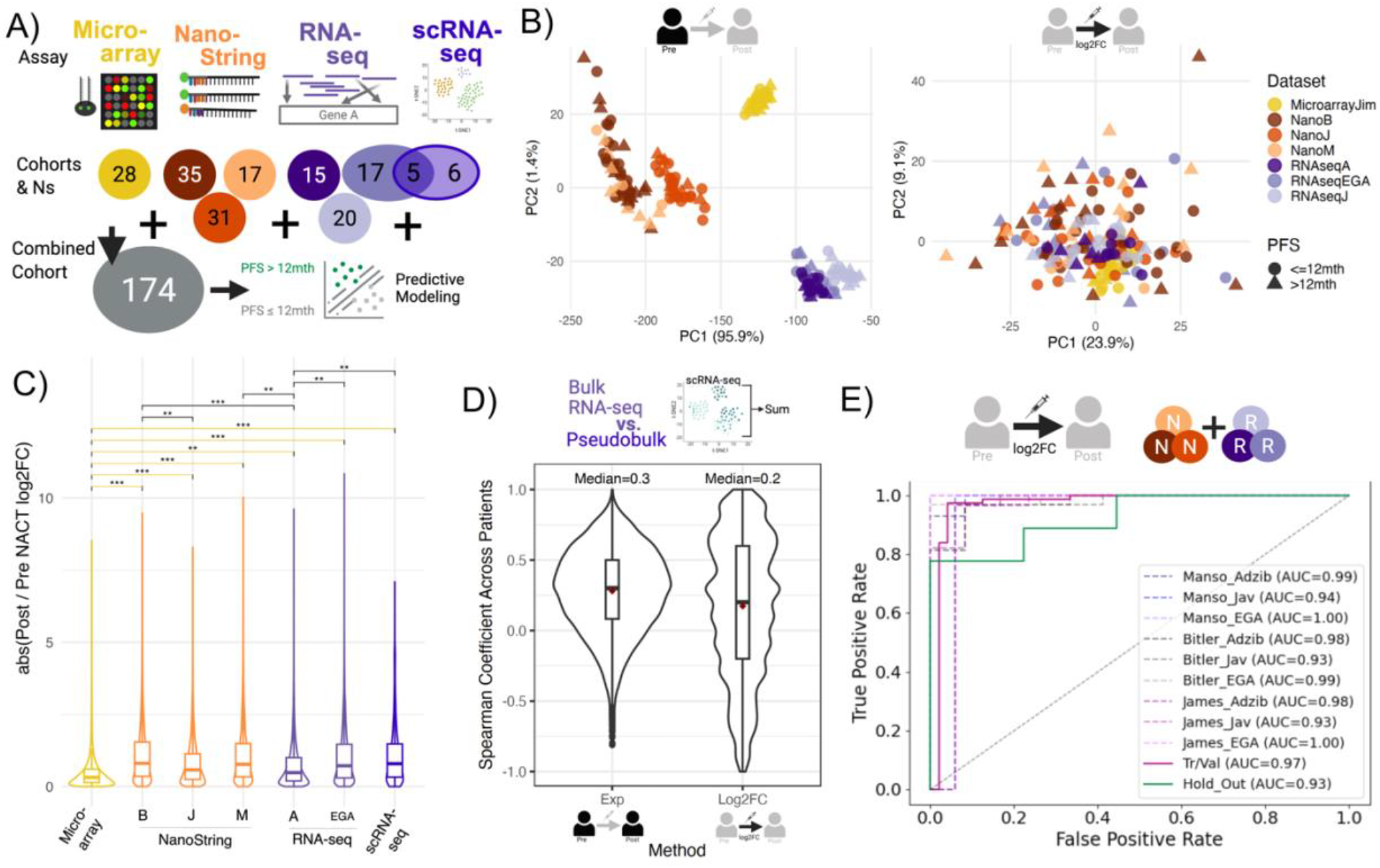
NanoString and RNA-seq show the lowest biases for direct combination of longitudinal cohorts but are still distinguishable. **A**. Outline of study where we assessed the most appropriate approach to combine longitudinal data across 4 transcriptomic assays of varying cohort sizes for predictive modeling of PFS. **B**. PCA across all patients/cohorts for the bulk assays using either normalized expression pre-NACT (left) or log2FC between normalized expression post and pre-NACT (right). **C**. Violin plot of absolute values of log2FC between normalized expression post and pre-NACT across the different assays/cohorts. Pairwise comparisons using Wilcoxon-Mann-Whitney tests and minimum effect sizes (Cliff’s delta) are shown with those between Microarray highlighted in yellow. Key: adjusted p-values below 0.01 (**) and 0.05 (*) with Cliff’s deltas ≥ 0.33, or adjusted p-values below 0.001 (***) and Cliff’s delta ≥ 0.474. **D**. Spearman correlation coefficient of gene expression across patients between normalized (CPM) expression levels pre or post NACT, or log2FC for bulk RNA-seq and pseudobulked scRNA-seq from the same FFPE samples. **E**. Receiver Operator Characteristic Curves of a model trained to predict NanoString samples using log2 Post/Pre NACT fold changes (AUC=Area under the curve) from the unmatched data set. Tr/Val = training/validation samples (N=128), Hold out (N=24), remaining cohorts are Tr/Val data of a NanoString mini-cohort combined with a RNA-seq cohort.

**Table 1.** Summary of Key Clinical Data of Longitudinal Cohorts. Med=Median. Full=Full data set, N+R=NanoString and RNA-seq data set combined. Some variables are not available for all patients. The number of patients with the data and their summary statistics are noted. Micro=Microarray. PFS = Platinum Free Survival. PFI = Platinum Free Interval. OS = Overall Survival. CRS=Chemo Resistance Score. PFI, PFS, and OS are in months. Articles for the different experiments are found in Methods.

|  | Full* | N+R | NanoString |  |  | RNA-seq |  |  | Micro | scRNA-seq |
| --- | --- | --- | --- | --- | --- | --- | --- | --- | --- | --- |
|  |  |  | Bitler | Manso | James | Adzib | Jav | EGA | Jim-Sanchez | Zhang |
| **PFS Median (N)** | **12**<br>(168*) | **11**<br>(140) | **9**<br>(35) | **18**<br>(17) | **10**<br>(31) | **11** (15) | **14.3**<br>(20) | **12.3**<br>(22) | **14**<br>(28) | **9.8**<br>(8) |
| **PFI Median (N)** | **6.8**<br>(29*) | --<br>(0) | --<br>(0) | --<br>(0) | --<br>(0) | --<br>(0) | --<br>(0) | **6.8**<br>(23) | --<br>(0) | **5.9**<br>(11) |
| **OS Median (N)** | **35**<br>(109*) | **33**<br>(81) | **34**<br>(35) | --<br>(0) | --<br>(0) | **36**<br>(6) | **22.9**<br>(20) | **30**<br>(21) | **40.5**<br>(28) | **30**<br>(8) |
| **Age Median (N)** | **63**<br>(136*) | **63.5**<br>(108) | **60**<br>(35) | --<br>(0) | **63**<br>(31) | **62.5**<br>(4) | **62**<br>(20) | **68**<br>(19) | **60**<br>(28) | **67**<br>(11) |
| **CRS Median (N)** | **2**<br>(51*) | **2**<br>(51) | --<br>(0) | **2**<br>(17) | --<br>(0) | --<br>(0) | **2**<br>(19) | **2**<br>(16) | --<br>(0) | **2**<br>(11) |
| **Stage N** | **101\*** | **73** | 0 | 0 | 31 | 0 | 20 | 23 | 28 | 11 |
| IIIA | 1* | 1 |  |  | 1 |  | 0 | 0 | 0 | 0 |
| IIIB | 3* | 3 |  |  | 3 |  | 0 | 0 | 0 | 0 |
| IIIC | 68* | 52 |  |  | 23 |  | 15 | 15 | 16 | 3 |
| IV | 16* | 4 |  |  | 4 |  | 0 | 0 | 12 | 0 |
| IVA | 6* | 6 |  |  | 0 |  | 1 | 5 | 0 | 7 |
| IVB | 7* | 7 |  |  | 0 |  | 4 | 3 | 0 | 1 |

\*scRNA-seq (Zhang) includes some patients also found in EGA so the Full Data set numbers account for only considering each patient once.

Overall, the combined transcriptomics cohort consisted of 174 patients with associated progression-free survival (PFS) or platinum-free interval (PFI) outcomes (Table 1, Supplemental Table 1). Although PFS and PFI are both commonly defined as the time interval between the completion of upfront chemotherapy and disease recurrence diagnosed via CA125 or imaging, the PFS[22,23] and PFI[14] of the 5 patients with bulk RNA-seq and scRNA-seq were not equivalent, even after considering timing differences of publications (Supplemental Table 1). Therefore, we considered PFS (in months) in all cases except for analyses confined to scRNA-seq alone. We then sought the most well-suited approaches to integrate data from the different assays to enable a single, combined cohort for predictive analysis of PFS (Figure 1A). This combined cohort where patients have assay-matched pre-post data will hereby be referred to as the “Single-assay pre-post cohort.” Similarly, a mini-cohort refers to the data from one study and assay (e.g. Bitler-NanoString, EGA-RNAseq), with the study name based on the first author or author with whom we mainly communicated.

### New double-assay post-NACT cohort

To directly interrogate assay biases of RNA-seq and NanoString, we constructed a new data set of 24 Formalin-Fixed Paraffin-Embedded (FFPE) HGSC post-NACT tumor samples from patients not previously assessed. Briefly, sequential FFPE tumor sections from an individual tumor block were used for either RNA-seq or NanoString. Henceforth, samples from this data set will be referred to as the “Double-assay cohort.”

### External microarray pre-NACT cohorts

Most HGSC transcriptomic data with associated recurrence/survival data are available as pre-chemotherapy microarray data. This includes two publicly available portals for Ovarian (CSIOVDB) or Serous Ovarian (KMPlot) cancer (which include The Cancer Genome Atlas (TCGA) data set)[24,25]. These data sets, however, are not specific to HGSC and do not easily allow cohort comparisons. Therefore, we also re-analyzed four pre-NACT microarray HGSC cohorts with overall survival information: Bonome (N=185)[26], Crijns (N=415)[27], Mok (N=53)[28], Yoshishara (N=260)[29].

## Modeling

### Predicted outcomes and modeling

We considered two outcomes to predict within the single-assay pre-post cohort. First, we trained a model to predict progression-free survival (PFS) categories based on the median of the cohort: PFS > 12 mths or ≤ 12 mths. Second, to characterize assay biases, we trained a separate model to predict whether a sample was sequenced with NanoString or RNA-seq. In all cases, to assess how robust the predictive power of a feature combination was to model parameters/structure, we considered three model architectures based on those most successful for HGSC-based modeling previously[15]: logistic regression using either LASSO or Ridge for regularization, and support vector machine. In all cases, because logistic regression performed as well or better than the support vector machine, we report logistic regression’s more interpretable coefficients. Area under the Receiver Operating Characteristic Curve (AUROC) was primarily used to compare model performance, including when using different gene numbers in cross-validation. Other model assessments are detailed in Supplementary Methods.

### Training, Validation, and Hold-out data

Before training a model, a “hold-out” data set was removed for final testing, defined as a minimum of 10 patients or 10% of the total cohort studied, equally representative of each classification, mini-cohort, and PFS levels (details in Supplemental Methods). Due to the small sample sizes, we used leave-one-out cross-validation (LOOCV) to determine the final number of genes to include in the model (k). The final model parameters were then estimated using the full training/validation data set.

### Feature Selection

Including the full feature (gene) list would likely lead to overfitting (due to small sample sizes) or convoluted comparison of performance across assays with different gene numbers. Therefore, we performed an initial feature selection to confine the feature space considered by a model to a maximum of 25 genes, as is often done in high-dimensional data sets[30]. When predicting PFS, genes were also first filtered based on the predicted limits of detection in expression and log2FC, as detailed in the Supplementary Methods.

Because the single-assay pre-post cohort comprised multiple mini-cohorts with differing sample sizes and technical contexts, we implemented two complementary feature selection strategies designed to mitigate cohort-specific biases. All strategies were implemented using training data only. The “Consensus” strategy considered genes shared among the top k predictive genes (measured by ANOVA-based F-scores) in each mini-cohort. The “Bootstrapping” strategy resampled from cohorts to estimate the underlying predictive value (measured by ANOVA-based F-scores) of genes. We bootstrapped samples from the training set (with equal number of patients within the two categories: either NanoString and RNA-seq or PFS > 12 mths and PFS ≤ 12 mths) using weights inversely proportional to the sample size of each mini-cohort. Therefore, assays or mini-cohorts with lower sample numbers were equally represented as those with higher sample numbers. With each bootstrapped sample, the 50 or 100 genes with top F-scores were recorded. For each gene, we calculated the cumulative proportion of bootstrapped samples that included it in this top-ranking list, a value hereby referred to as predictive frequency. The genes with the highest-ranked predictive frequencies from bootstrapping or consensus strategies were then considered for LOOCV to determine the final gene number in the model (details in Supplemental Methods).

## Data Availability

For NanoString, we reanalyzed the raw files found in GEO from GSE319500[1], [15] (Bitler), GSE181597[18] (Manso), and GSE201600[20] (James). For RNA-seq, the Adzib data set used raw files from GEO GSE227100[13]. Survival information was provided for a subset of patients by the original authors. The EGA data set was so named since raw files could only be collected from the European Genome Archive (EGAD00001006456). This data set included consortia of HGSC studies MUPETFaasi2, HERCULES, and CHEMORESPONSE. The metadata and previous analyses of the data were collected from [22,23,31,32]. The Jav data set only provided normalized counts (RPKM) as the publication’s Supplemental Table 5[21]. For microarrays (Jim-Sanchez), normalized data were found at GSE146963[16]. Non-pre-post microarray data was downloaded from GEO numbers GSE32062 (Yoshishara)[29], GSE26712 (Bonome)[26], GSE13876 (Crijns)[27], and GSE18521 (Mok)[28]. Quality Control counts and metadata for scRNA-seq (Zhang) data from [14] were downloaded from GEO using accession number GSE165897 (GSE165897_cellInfo_HGSC.tsv and GSE165897_UMIcounts_HGSC.tsv.gz). Data from the double-assay cohort is at GSE323347 (RNA-seq) and GSE322784 (NanoString). TOM similarity data can be found on zenodo at 10.5281/zenodo.19196835.

Users can also explore the data on https://hopetownsend.shinyapps.io/HGSOC-gene-explorer/. This app runs through Shiny apps free portal and therefore users are also encouraged to follow instructions on the Github repository (https://github.com/Hope2925/NanoString-RNAseq-HGSOC) to run it locally to avoid server limitations. The same repository contains all code, data, and results for this work.

## Results

### NanoString and RNA-seq show the greatest potential for direct combination of longitudinal data

#### Pre-post data does not address microarray and scRNA-seq biases

Technical biases of assays may be addressable through preprocessing, or may instead reflect fundamental and unalterable differences in data (e.g., from assays measuring biologically different features or having different detection/oversaturation limits). For example, within-assay scaling is often used to align assay expression data but assumes that all assays measure similar dynamic ranges (i.e., the lowest and highest values in each assay should be biologically comparable). Longitudinal-based calculations (like Post vs. Pre NACT log2 fold change (log2FC)) would address scale but not fundamental differences between data.

We therefore assessed the strengths and weaknesses of each assay, clarifying whether such differences could likely be addressed by preprocessing steps or were so fundamental as to hinder combinations for predictive modeling. To qualify the general effects of assay, and whether log2FC could limit biases from assay measurements, we considered how assay data aligned in low-dimensional space when using expression or Post vs. Pre NACT log2FC, based on Principal Component Analysis (PCA) (details in Methods). The shared 731 genes were used in this analysis (in NanoString and microarray probe sets and the GRCh38.p14 RefSeq annotation), with limitations of a shared feature space noted in Supplemental Results. We then directly assessed if using log2FC might improve the historically weak correlation of expression levels for bulk RNA-seq and scRNA-seq pseudobulk[33], by leveraging the fact that some patients were sequenced with both scRNA-seq and bulk RNA-seq.

Patients did cluster based on assay after dimensionality reduction from pre- or post-expression levels alone. However, using log2FC between the post and pre samples reduced this difference, making the assays and cohorts substantially overlap in low-dimensional space (Figure 1B). Within-assay scaling further improved overlap (Figure 1C). We note that, as found in previous work[8], microarray data showed evidence of lower dynamic range: lower log2FCs and interquartile ranges of expression compared to all other assays and cohorts (all adjusted p-values < 0.01 and effect sizes (Cliff’s deltas) > 0.33; Figure 1C, Supplemental Figure S1A and B). Thus, even under log2FC measures, microarray data contributed little variance across the combined cohort unless using within-assay scaling (Supplemental Figure S1C). These patterns imply that within-assay scaling can seem to harmonize data but does so by assuming comparable underlying dynamic ranges, which is likely violated for microarray data. In subsequent analyses, we use unscaled log2FC, and we consider microarray directionality to verify rather than train our models.

When comparing pseudobulked scRNA-seq and bulk RNA-seq, we observed the expected poor correlation using expression levels (N=13), yet saw similar trends in pre-post log2FCs (N=5) (median spearman coefficients (*ρ*) of 0.3 and 0.2, respectively) (Figure 1D). The resolution of scRNA-seq means that only the most highly expressed genes have sufficient read coverage for confident downstream analyses[33]. Although we hypothesized this poor correlation was from genes with low expression, the trends were common across all levels of expression (Supplemental Figure S2). Ultimately, we observed biases from pseudobulking scRNA-seq that limited its potential direct combination with bulk data sets.

#### NanoString and RNA-seq have limited biases influencing clinical predictive modeling

To assess if there are addressable biases between NanoString and RNA-seq, we tested whether a predictive model could distinguish between samples sequenced with RNA-seq or NanoString, using Post/Pre-NACT log2FC alone. We also assessed whether assay biases might impact clinical modeling by checking whether genes predictive of assay differences were also predictive of PFS. Specifically, we compared the predictive frequency of genes when predicting NanoString vs RNA-seq or PFS category, i.e., the proportion of bootstrapped samples where each gene was one of the top 100-ranked predictive features.

First, we trained a model that consistently predicted whether a patient was assayed by NanoString or RNA-seq, regardless of mini-cohorts considered (hold-out AUROC=0.93; Figure 1E; details in Methods). Nine of the twelve genes used by the final model (including the eight with the highest coefficients) showed the same directional differences between assays across all six mini-cohorts (Supplemental Figure S3). Therefore, several genes appear to have consistently different measurements between NanoString and RNA-seq. We also found evidence that RNA-seq and NanoString had poor concordance of predictive features, which may impact clinical modeling. The correlation of predictive frequencies of genes for PFS was low between assays (*ρ* = 0.028). Indeed, this correlation between assays was lower than the correlation of the gene predictive frequencies when distinguishing PFS status compared to NanoString from RNA-seq (*ρ* = 0.071, *ρ* = 0.077) (Supplemental Figure S4). These findings indicate generally high heterogeneity between RNA-seq and NanoString, and that genes relevant to PFS also have distinguishable differences between NanoString and RNA-seq. Indeed, several genes showing predictive frequencies of PFS above 0.3 in RNA-seq or NanoString cohorts also showed predictive frequencies above 0.3 when distinguishing NanoString and RNA-seq samples (Supplemental Figure S4). These findings suggest that calculating log2FC in longitudinal data is insufficient to remove assay-based biases in data, however, such heterogeneity across assays might be explained by unoptimized preprocessing steps of NanoString and RNA-seq.

### Differences between NanoString and RNA-seq measurements are largely addressable with data processing

Because NanoString and RNA-seq showed comparable dynamic ranges in our samples, their data could potentially be combined for predictive modeling by optimizing their respective preprocessing. To test this hypothesis, we considered our new double-assay cohort of 24 FFPE HGSC tumor samples, in which sequential FFPE tumor sections were processed using either RNA-seq or NanoString (Figure 2A). We evaluated the impact of two preprocessing components: 1) removing genes lowly expressed based on limits of detection, and 2) harmonizing regions considered in RNA-seq. Normalization approaches had little to no impact (Supplemental Results).

**Figure 2.**
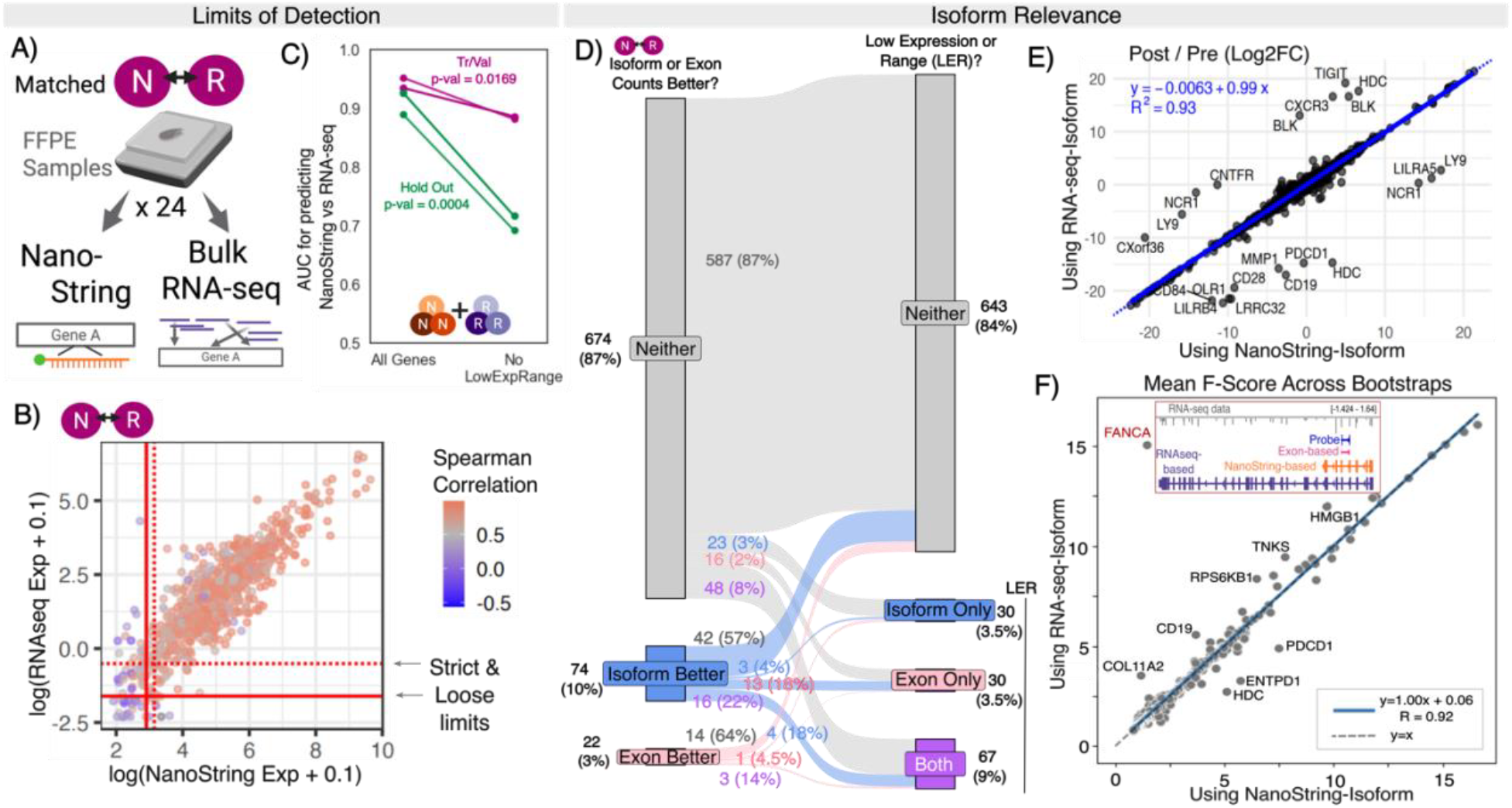
NanoString and RNA-seq assay-specific biases are largely due to limits of detection. **A**. Sequential samples from 24 FFPE isolations were sequenced with either NanoString or RNA-seq. **B**. Scatter plot of logged median expression from RNA-seq (Isoform-RPKM, y-axis) and NanoString (x-axis) for each shared gene (N=770). Colored by correlation between the double-assay data sets. Strict and loose limits of detection are labeled. **C**. AUROC of models predicting NanoString vs RNA-seq when allowing consideration of all NanoString genes vs those above the limits of detection for range and expression (No LowExpRange). **D**. Flow graph of the number of genes where isoform- or exon-based counting improves harmonization, and second whether the isoform or exon-based counts are below the limits of detection. **E**. Scatterplot of log2FC values of genes using RNA-seq based isoforms (y-axis) vs NanoString based isoforms (x-axis). Genes with differences ≥ 5 are labeled. **F**. Scatter plot of mean F-score across bootstrapped samples of genes when using RNA-seq based isoforms (y-axis) vs NanoString based isoforms (x-axis). The inset shows the probe-, exon-, NanoString isoform-, and RNA-seq isoform-based coordinates for *FANCA*.

If a gene is not sufficiently well expressed to be accurately detected, or if it has a narrow expression range, there may be insufficient variance in the measured levels to accurately estimate covariance, and therefore correlation. Therefore, we estimated the limits of detection of both expression and range for RNA-seq and NanoString harmonization by characterizing the levels and ranges, at which we observed a clear (strict) or starting (loose) decline in correlation between RNA-seq and NanoString (details in Supplemental Methods; final cutoffs in Supplemental Table 2; Figure 2B). An obvious difference between the assays is that while NanoString commonly only considers < 200bp of a gene (via probe), RNA-seq can consider any gene sequence(Figure 1A, Supplemental Figure S5A). Therefore, we evaluated if harmonization was impacted by counting across isoforms versus exons most closely corresponding to the NanoString probe, with the latter expected to resemble probe regions (Supplemental Figure S5). These results can be explored in our web portal (Data Availability Statement).

#### Most consistent differences are due to limits of detection

Projection of the double-assay cohort into a low-dimensional space suggested that although absolute values differed between RNA-seq and NanoString, relative gene expression across patients and dynamic range were maintained (Supplemental Figure S6). Regardless of using different normalizations or isoform-vs exon-based counting, most (51%) genes showed Spearman correlations above 0.85, with 17% consistently showing below 0.85. The remaining genes (32%) showed correlations that depended on preprocessing approaches (details in Methods).

Because counting differences had a limited impact on the harmonization of most genes, most poor correlations were likely from technical limitations of detection. Indeed, Spearman correlation coefficients were significantly lower in genes with the lowest expression and/or range quantile (adjusted p-values < 0.001, Supplemental Figure S7). When using cutoffs based on our strict predicted limits of detection, only 31 genes still had consistent correlation coefficients below 0.7. *TMEM140, RAD50, PVRIG, HDAC3, BAD*, and *ELOB* had coefficients far below 0.5, possibly partly from low expression (< 5 TPM). Removing genes below the limits of detection also lessened the distinguishability of NanoString and RNA-seq samples using log2FC (hold-out AUROC from 0.93 to 0.65) (Figure 2C). In this case, the model relied more on overfitting to noise across cohorts rather than assay-specific differences, with only two of the twelve model genes exhibiting consistent trends (Supplemental Figure S8). Ten of the twelve genes included in the unrestricted model we trained for distinguishing NanoString and RNA-seq samples fell below these limits of detection, further supporting the conclusion that differences in detection limits explain most systematic differences between assays (Supplemental Figure S3).

#### Probe-focused counting can harm rather than improve harmonization

Remaining systematic differences might be resolved with an RNA-seq counting approach. A total of 96 genes (13%) had improved Spearman correlation coefficients between NanoString and RNA-seq when using either exon- or isoform-based counts (increase of > 0.05 in at least two of three normalization approaches) (Figure 2D left). Exon-based counts may improve region matching between assays, but also decrease data by considering a smaller genomic segment. Indeed, we found that isoform-based counts commonly performed better due to simply increasing data above the limits of detection: 40% of genes for which isoform-based counts were better had expression levels below detection limits (Figure 2D Isoform-Better to Exon Only and Both). Similarly, genes with better harmonization from exon-based counts had higher exon-but not isoform-based counts than genes with better isoform-based correlation (adjusted p-value < 1*x*10^−26^; Supplemental Figure S9). Still, most cases where exon- or isoform-based counting was preferential (64% and 57%, respectively) were not explained by detection limits (Figure 2D), and probe-focused counting had limited, if any, improvement in harmonization (Supplemental Results; Supplemental Figures S10, S11, S12).

Clinically relevant insights could also be lost by restricting RNA-seq analysis to isoforms best matching NanoString probes rather than the most highly expressed ones. However, there was high correlation (*R*^2^ =0.93) of log2FC in genes when using the highest-expressed or NanoString isoform, even though 59% of genes had a highest-expressed isoform different from the NanoString one (Figure 2E). All outliers (> 5 change in log2FC; labeled in Figure 2E) were explained by expression levels changing between zero and values already below detection limits (details in Supplemental Results). Isoform usage also did not impact NanoString and RNA-seq harmonization; in our double-assay data set, despite 462/770 genes (60%) having a highest-expressed isoform different from the NanoString one, we observed little to no change in Spearman correlation between NanoString and RNA-seq expression levels (Supplemental Figure S13A). Finally, isoform-based changes in log2FC lead to minimal differences in genes’ predictive power of NanoString vs RNA-seq samples or PFS (as measured by mean F-score across bootstraps) (Figure 2F and Supplemental Figure S13B, *R*^2^=0.92 and *R*^2^=0.99). The only obvious outlier was *FANCA* whose isoform changed from the shortest to the longest form (increased length by more than 50kb) (subset of Figure 2F). Therefore, isoforms used made minor impact on predictive modeling except with extreme region changes (e.g. more than 50kb). Overall, preprocessing steps focused on normalization and counting approach had limited impacts on harmonization across NanoString and RNA-seq.

### Integrating NanoString and RNA-seq pre-post longitudinal data allows generalizable survival prediction

NanoString and RNA-seq showed enough congruence after preprocessing to be combined into a single cohort for predicting PFS with the shared gene space (N=770) (Figure 3A). The extent to which such a combination improves modeling has not been previously determined. Therefore, we trained a PFS predictive model on a “combined cohort” of the single-assay pre-post NanoString and RNA-seq cohorts, using scRNA-seq and microarray data for external verification. Since scRNA-seq had 5 individuals whose bulk RNA-seq were used for model training, we could consider external verification with both assay type and people. As noted previously, because microarray data exhibits a lower dynamic range, we used microarray data only to validate whether the pre-post change direction agreed with the predictions of our model. Similarly, scRNA-seq could clarify which cell-states best represent a given gene’s bulk signal. Finally, we considered the source for the vast majority of HGSC data, pre-NACT microarray, to evaluate if our log2FC-based biomarkers showed association with survival metrics based on microarray pre-NACT expression alone. Briefly, results showing how our harmonization insights improve upon previous single-assay predictive modeling of PFS are in Supplemental Results. The predictivity of these genes and similar analyses can be explored in our web portal (Data Availability).

**Figure 3.**
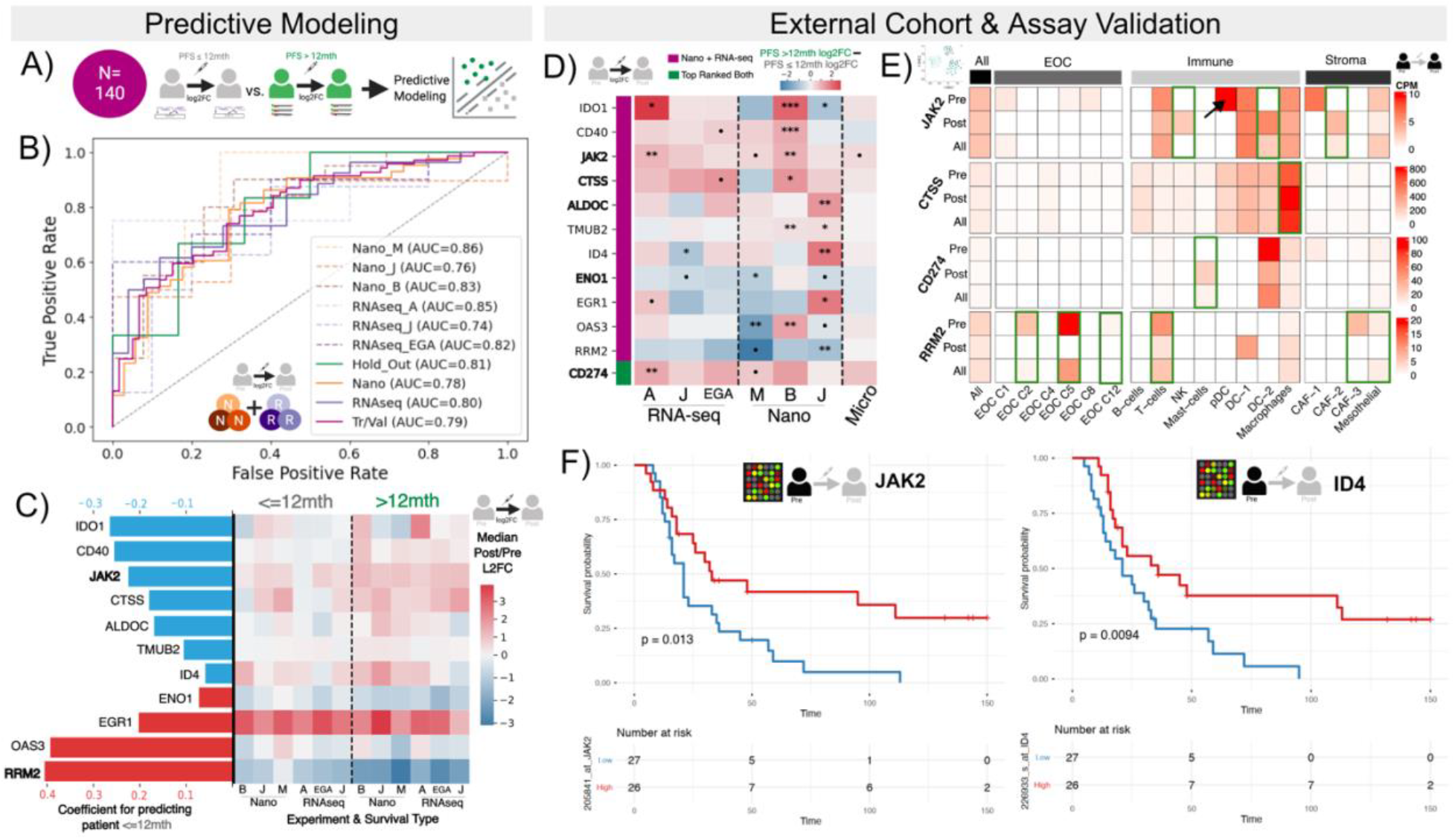
Predictive modeling in combined cohort of NanoString and RNA-seq reveals consistent biomarkers. **A**. Combined cohort of NanoString and RNA-seq used to perform predictive modeling of PFS. **B**. Receiver Operator Characteristic Curves of a model trained to predict if a patient has a PFS≤ 12 mths using log2 Post/Pre NACT fold changes. Tr/Val = training/validation samples (N=128), Hold Out N=12. **C**. Coefficient values of model genes where (blue indicates a negative and red positive coefficient for the log2FC). Bolded genes have consistent trends across all 6 mini-cohorts. Heatmap of median log2FC in each mini-cohort split by PFS category. **D**. Difference between median Log2FC of patients with PFS > 12 mths and PFS ≤ 12 mths of combined model genes across the 7 mini-cohorts with the p-values of t-tests of the log2FC dictated by •< 0.1,* < 0.05,** < 0.01,*** < 0.001. **E**. Heatmap of counts per million (CPM) per pseudobulked cell-state from single-cell data pre-NACT, post-NACT, and combined (All). **F**. Kaplan-Meier curves based on median expression split of *JAK2* and *ID4* in an external pre-NACT Microarray cohort.

A model that used log2FC of genes to classify patients based on the median PFS (< 12 mths) achieved AUROC > 0.7 (0.81 for hold-out) across all mini-cohorts and assays (Figure 3B). Further results confirming the model learned biological patterns and feature relevance are provided in Supplemental Results (Supplemental Figures S15, S16A, S17). All eleven combined model genes except *TMUB2* had explicit literature supporting their predictive value for HGSC survival or differential expression in HGSC cells with varying chemotherapeutic resistance (Supplemental Table 3 and Results). Seven had more than one source supporting their linkage: *IDO1, CD40, JAK2, ID4, ENO1, EGR1*, and *RRM2*. Despite the external validation of most model genes in HGSC, only *JAK2* and *RRM2* showed consistent trends in PFS > 12 mths and PFS ≤ 12 mths across all mini-cohorts: stronger median upregulation and downregulation with PFS > 12 mths patients of *JAK2* and *RRM2*, respectively (Figure 3C). Based on the DepMap database[34] of CRISPR knockouts and RNA knockdowns, most cancer cell lines showed a high dependence on *RRM2* (Supplemental Figure S18). OVCAR8, an HGSC cell line characterized by high cisplatin resistance, was classified as *RRM2*-essential by DepMap cutoffs.

#### Verifying results in external scRNA-seq and microarray

For external verification in scRNA-seq and pre-NACT microarray data, we considered 12 genes: the 11 combined model genes, and *CD274* [protein PD-L1]. *CD274* was included because it was among the highest and consistently top-ranked genes in our feature selection for both RNA-seq and NanoString, and was also a top marker in a previous PFS predictive model for HGSC[15] (details in Supplemental Results; Supplemental Figure S19). Of the 12 genes, 5 (42%) showed the same directionality across NanoString, RNA-seq, and microarray, with a maximum of one mini-cohort within NanoString or RNA-seq disagreeing (Figure 3D). In some cases, the microarray data’s log2FC directionality clarified an initial disagreement between mini-cohorts, e.g., supporting a finding that higher upregulation of *IDO1* generally corresponds to PFS > 12 mths.

scRNA-seq analysis indicated the cell types through which the four genes with the most consistent signals were likely conferring impact on PFS (results for all in Github repository). While *JAK2* showed the strongest initial signal in plasmacytoid dendritic (pDC) cells, it showed increased expression post-NACT in NK and dendritic cells (DC-2) (Figure 3E). *CTSS* and *CD274* showed increased post-NACT expression in macrophages and mast cells, respectively. Interestingly, the scRNA-seq data showed the opposite correlational direction of PFI with *CD274* to bulk, indicating that decreased post-NACT expression correlated with higher PFI (Supplemental Figure S20A). As noted previously, PFI was used here since not all samples had PFS. Finally, *RRM2* showed the clearest decrease in Epithelial Ovarian Cancer (EOC) Cluster 5, which had previously been annotated as proliferative cancer cells (Figure 3E)[14]. This finding corresponds with a stronger decline of *RRM2* expression reflecting lower proliferation of cancer cells and therefore serving as a biomarker of treatment success as suggested by our model. *EGR1* and *IDO1* had log2FC significantly correlated (Spearman correlation p-value below 0.05) with the PFI in cancer-associated fibroblasts, but the correlational signal was dominated only by patients already used for model training (Supplemental Figure S20A and B).

Finally, all 12 genes were significantly associated with survival and/or PFS in Ovarian (CSIOVDB)[24] or Serous Ovarian (KMPlot)[25] cancer (Supplemental Table 3). When considering HGSC-specific pre-NACT microarray data, *JAK2* and *ID4* showed significant association (adjusted p-value < 0.05; univariate Cox regression) with survival across multiple cohorts in the same direction, including the cohort with the smallest N (Mok, N=53; Hazards Ratio 95% confidence intervals of [0.23-0.85] and [0.22-0.83]; p-values of 0.01) (Figure 3F). Seven model genes (58%) showed significant association in at least one of these four HGSC cohorts (Supplemental Table 4): *IDO1, JAK2, CTSS, ID4, EGR1, OAS3, CD274*.

Overall, these findings indicate that data from NanoString and RNA-seq can be effectively combined to identify biomarkers, after optimizing preprocessing steps.

### Full genome consideration reveals key genes missing and provides additional mechanistic context

#### Non-NanoString panel genes mark treatment success

We evaluated the unique insights that assays considering all genes (RNA-seq and scRNA-seq) could provide for discovery. To this end, we trained a model for predicting patients with PFS ≤ 12 mths, using the 57 patients with matched pre-post RNA-seq and all genes transcribed. Importantly, our feature selection approach ensured that there was no obvious burden from including more genes (details in Methods).

This model achieved higher AUROCs when trained on just the RNA-seq data than we observed with the combined (NanoString + RNA-seq) model (0.86 vs original 0.81 in hold-out set) (Figure 4A). Further results validating the RNA-seq model and feature importance are in Supplemental Results (Supplemental Figures S14, S16B, S22). Six of the nine RNA-seq model genes showed consistent median log2FC differences between PFS classes across mini-cohorts (highlighted in 4B). This is a higher number than observed with the combined model, but we also only needed three instead of six mini-cohorts to demonstrate similarity. Seven of the nine RNA-seq model genes also had literature support linking them specifically to ovarian cancer or chemoresistance: *GBP4, AMOTL2, ESTY3, HCN3, DDX11, CLK2*, and *CDKL2* (Supplemental Table 3 and Results). Although *GBP4* was the only RNA-seq model gene in the NanoString panel, NanoString panel genes generally showed higher predictive power (Mean F-score across bootstrapped samples) than other genes (p-value < 0.001, Supplemental Figure S21). Specifically, *CD274* and *GBP4* were consistently within the top 25 genes with the highest predictive frequency when only considering NanoString panel genes across the combined cohort.

**Figure 4.**
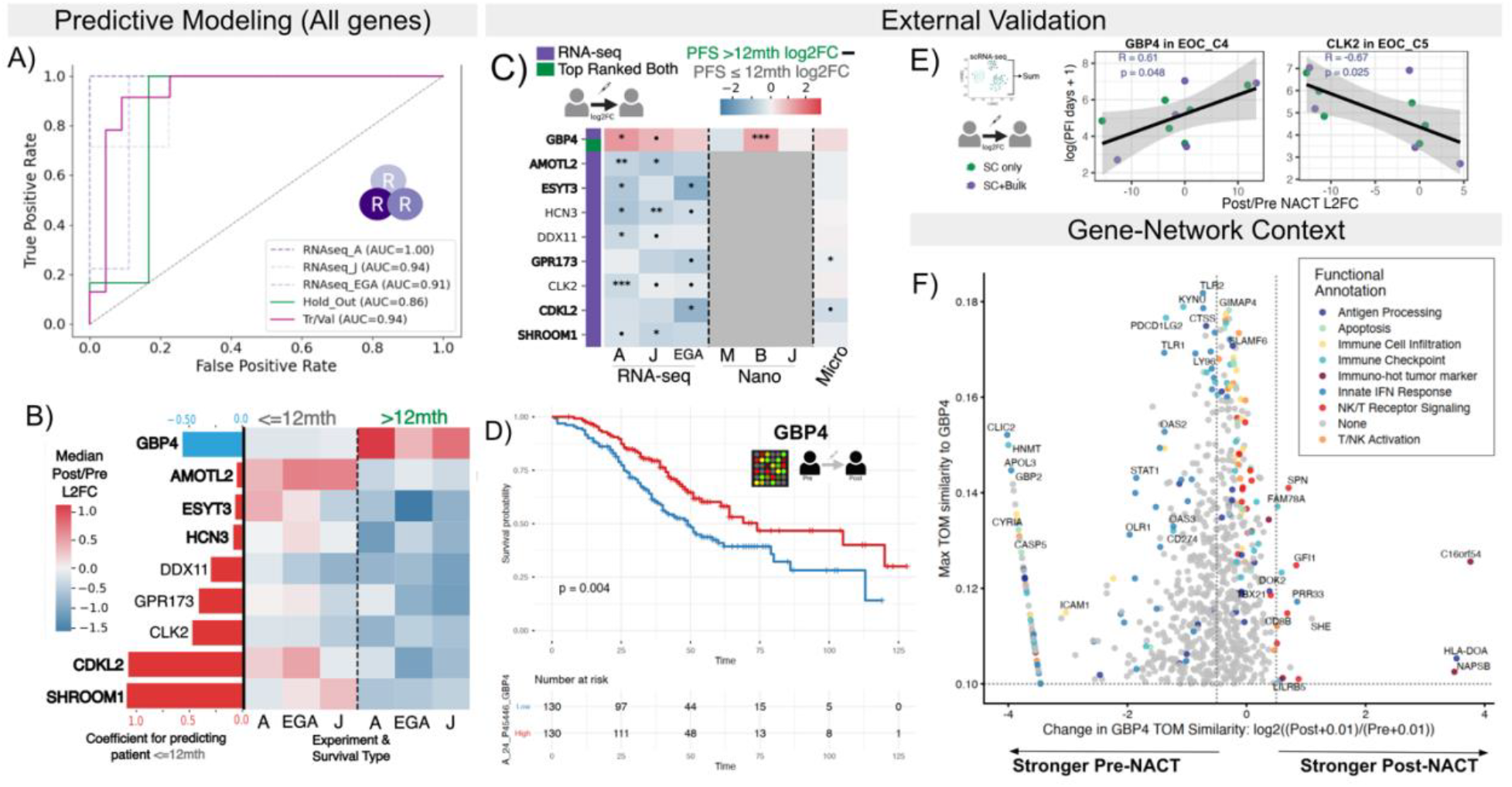
Full genome analysis improves mechanistic understanding of biomarkers. **A**. Receiver Operator Characteristic Curves of a model trained to predict if a patient has a PFS≤ 12*months* using log2 Post/Pre NACT fold changes using all RNA-seq genes and data. Tr/Val = training/validation samples (N=45), Hold Out N=12. **B**. Coefficient values for each of RNA-seq model genes (blue= negative and red=positive coefficient for log2FC). Bolded genes have consistent trends across all 3 mini-cohorts. Heatmap of median log2FC in each mini-cohort of patients split by PFS category. **C**. Difference between median Log2FC of patients with PFS > 12 mths and PFS ≤ 12 mths of RNA-seq model genes across the mini-cohorts, with the p-values of t-tests of the log2FC dictated by •< 0.1,* < 0.05,** < 0.01,*** < 0.001. **D**. Kaplan-Meier curve based on median expression split of *GBP4* in an external Microarray cohort. **E**. Scatter plots of Post/Pre NACT Log2FC from pseudobulked scRNA-seq data (N=11, within cell-type) and the log(PFI + 1). Fitted line with R and p-values from a linear regression with standard error highlighted. **F**. Volcano-like plot of genes’ changes in TOM similarity with GBP4 in Pre- and Post-NACT WGCNA networks (x-axis) and the maximum similarity (y-axis). Genes determined as shared or post/pre only are colored based on general immune-based functions, with genes of highly mixed or lesser known functions labeled as None.

As with the combined model, microarray and scRNA-seq data supported RNA-seq model findings. Six of the nine genes (67%) showed the same log2FC directionality associated with PFS across all available assays, with a maximum of one NanoString or RNA-seq mini-cohort showing disagreement (Figure 4C). Similarly, all genes showed pre-NACT expression significantly associated with overall survival and/or PFS in Ovarian (CSIOVDB)[24] or Serous Ovarian (KMPlot)[25] cancer (Supplemental Table 3). Pre-NACT expression of only three of the model genes (33%, *GBP4, DDX11*, and *CLK2*) was significantly associated with survival (adjusted p-value < 0.05 univariate Cox regression) in at least one of the four HGSC-specific microarray cohorts (Supplemental Table 4). *GBP4* was significant across multiple cohorts in the same direction, with a Kaplan-Meier curve of the Japanese Yoshihara cohort[29] shown in Figure 4D (Hazards Ratio 95% confidence interval [0.41-0.85], p-value=0.004). *GBP4* showed the most consistent results among all guanylate-binding genes in our and other data sets (Supplemental Results; Supplemental Figure S23). Both *GBP4* and *CLK2* also had log2FC correlated with PFI in cancer cells according to scRNA-seq, in the same directions suggested by the bulk RNA-seq model. *GBP4* showed upregulation in patients with longer PFI and downregulation in those with shorter PFIs in the proliferative cancer cells (EOC C5); stronger downregulation of *CLK2* was associated with higher PFI in dendritic cells and EOC C4 (annotated as differentiating cancer cells) (Figure 4E, Supplemental Figure S20)[14]. Log2FCs of four additional model genes (total 67%) also correlated with PFI: *HCN3, AMOTL2, ESYT3, GPR173* (Supplemental Figure S20). Overall, being reduced to a gene panel list, even one specific to cancer, can hinder discovery of relevant biomarkers.

#### Gene networks clarify mechanistic relevance of *GBP4*

Considering a full gene space also allowed us to perform robust network analyses to hypothesize 1) which genes/pathways a gene correlated with and therefore might be represented by that gene in the model, and 2) the contextual role of a gene itself. We focused on *GBP4* due to its differential signal being highly consistent. Briefly, we reconstituted pre- or post-NACT networks using the topological overlap matrix (TOM) similarity scores (measurement of gene interconnectedness) between genes from WGCNA[35] (RNA-seq) and hdWGCNA[36] (scRNA-seq) (details in Supplemental Methods).

We first evaluated how the bulk-based gene-networks of *GBP4* changed based on treatment status (Pre-vs Post-NACT) (details in Methods). Regardless of timing, *GBP4* connected to a core set of immune-relevant genes including immune checkpoints, lymphocyte regulation, chemokine signaling, and immune-cell infiltration (Figure 4F, Supplemental Table 5). This finding supports that *GBP4* indeed corresponds to an immune role in the HGSC context. In Pre-NACT networks, *GBP4* was connected to genes primarily involved in innate immunity: innate immune sensing, apoptosis, antigen processing, interferon-stimulated genes, metabolic stress, and immune inhibitory checkpoints (including *CD274* [PD-L1] and *PDCD1LG2* [PD-L2])) (Figure 4F, Supplemental Table 5). Enriched gene-ontology pathways (q value < 0.05) unique to the pre-NACT gene group included viral response, interferon-mediated signaling pathways, and positive regulation of Tumor Necrosis Factor. Conversely, post-NACT *GBP4* connectivity shifted closer to pathways of cytotoxic lymphocytes, with enriched pathways (q value < 0.05) unique to the post-NACT group including thymic T cell selection, T helper (CD4+) cell differentiation and immune response, and regulation of alpha-beta T-cell differentiation. Post-specific genes directed towards T helper cell fates and expansion, marked cytotoxic T-cell proliferation, increased CD4+ and CD8+ T cell trafficking, and activated T/NK cells (Figure 4F, Supplemental Table 5). *HLA-DOA* and *CLEC10A* both mediate antigen loading in macrophages and dendritic cells, thereby influencing T helper responses. Finally, *PTPN7, NAPSB, C16orf54*, and *TMC8* have been independently associated with immune-hot tumor microenvironments and serve as predictive markers of cancer prognosis[37-40] (Figure 4F). Overall, these findings indicate that *GBP4* networking may mark a transition from a general innate response to one of cytotoxicity.

Single-cell expression and networks support this interpretation. *GBP4*-associated edge weights from bulk RNA-seq most closely matched the scRNA-seq networks of the dendritic cells in which *GBP4* was also most strongly expressed prior to treatment (Supplemental Figure S24A,B). Following NACT, *GBP4* expression then decreased markedly in macrophages, NK cells, B cells, and dendritic cells, while remaining stable in T cells and increasing in innate lymphoid cells and mast cells (Supplemental Figure S24B). *GBP4* network correspondence also shifted from primarily dendritic cells to include the NK cell group (NK+ILC, with 86% NK cells), with the caveat that NK cell counts above 50 were observed in only two patients pre-NACT. Together, these results indicate that *GBP4* expression and network context are both reprogrammed by chemotherapy, so that its expression patterns may reflect a shift from a myeloid-dominated, immunosuppressive environment toward lymphoid-associated cytotoxic immune programs in our model.

## Discussion

### Foundational guidance on data integration across assays for predictive modeling

In this study, we evaluated the degree to which distinct transcriptomic technologies can be integrated, and provided guidelines for combining assay data in predictive modeling and validation. After identifying the strengths and weaknesses of each platform, we established harmonization procedures for NanoString and RNA-seq, which showed sufficient consistency for direct comparison even in variable sample types like FFPE. Predictive modeling and network analysis further demonstrated how integrating probe-based assays with approaches that consider full gene space and single-cell dynamics can identify biomarkers and their mechanistic context.

Our work indicated that microarray and scRNA-seq contain systematic biases that should be carefully considered before integration with NanoString or RNA-seq data. Although percentile scaling is commonly used to incorporate microarray data sets, both our and previous work suggests that this approach is misleading because microarrays do not share dynamic ranges with other assays[8]. A more robust approach may be to incorporate a binary microarray indicator in models trained across bulk assays. Matched pre-post data also did not resolve poor correlations between pseudobulk scRNA-seq and bulk RNA-seq, although limited sample size restricted our ability to assess the full extent and reasons for this divergence. Prior work showed that smaller, robust cells like immune cells are preferentially captured in scRNA-seq[41], potentially altering cell-type proportions and supporting the use of RNA-seq deconvolution harmonization approaches instead.

Genes with low expression are a specific and substantial barrier to assay integration. Determining the limits of detection for single and multi-assay studies is thus an essential step prior to combining data from different assays. Our analyses further suggested that typical RNA-seq depth rarely allows region-specific counting to influence NanoString–RNA-seq harmonization except for large (> 50 kb) isoform changes. Our web portal enables exploration of gene-level harmonization using isoform- and exon-based counts.

Finally, we propose guidelines for integrating transcriptomic data sets as technologies evolve. While NanoString panels are gene-limited, they provide an efficient and often cheaper alternative to RNA-seq. Meanwhile, RNA-seq’s extensive feature set provides a promising way to identify the relevant sub-1000 gene sets for NanoString panels suitable for clinical adoption. NanoString and RNA-seq data sets can then be combined for improved predictive modeling, while RNA-seq and scRNA-seq help characterize mechanistic context underlying biomarkers. For example, we clarified that *GBP4* is not necessarily an effector itself, but marks shifts in the tumor microenvironment from immune suppression and generalized inflammatory towards specific cytotoxic, immune-hot tumor signatures.

### Potential biomarkers and models for downstream experimentation in HGSC

These guidelines enabled the discovery of predictive models and biomarkers of treatment response in HGSC that can guide hypothesis generation and focused experimental work.

Our findings suggested that high expression of *RRM2* reflected the proliferation of cancerous cells and, therefore, chemoresistance. Indeed, inhibition of ribonucleotide reductase (RNR), whose activity RRM2 regulates, increased the sensitivity of chemo-resistant cancers in multiple clinical trials, including one in platinum-resistant ovarian cancer[42], [43]. However, direct RNR inhibition can lead to severe side effects, motivating efforts to develop expression-level inhibition of *RRM2* instead[42].

Interestingly, our data suggested that increased expression of *CD274* [PD-L1], and low initial expression and high post-NACT expression, correlate with increased PFS. However, PD-L1 binding inhibits T-cell activation and cytokine production, making it a key therapeutic target[44]. We propose two explanations for this apparent contradiction. First, increased *CD274* expression may reflect both a potential immunosuppression and a responding immune system, as NF-kB induces *CD274* expression[44]. Supporting this hypothesis, we observed that post-NACT *CD274* expression occurred primarily in immune rather than cancer cells, and lower expression within immune cell types correlated with higher PFS. Second, increased bulk *CD274* expression may indicate an increased proportion of immune cells in the samples.

Finally, increased and generally high post-NACT expression of *JAK2* and *GBP4* correlated with higher PFS. Although functional assays in cancer cell lines have suggested that decreased *JAK2* expression limits proliferation[45], [46], our findings indicated that *JAK2* expression in HGSC tumors occurs primarily in immune rather than epithelial cells. This indicates the importance of considering a gene within the full tumor microenvironment. Indeed, JAK inhibitors only slightly improved chemosensitivity in mice with tumors already exhibiting elevated expression of JAK genes[46]. While a useful biomarker, JAK2’s diverse cellular roles may limit its therapeutic potential. Instead, *GBP4* shows broader expression across cell types and has been described as a marker of immune-hot tumors[47]. Experimental studies have indicated *GBP5* and *GBP1* expression are critical to preventing ovarian cancer proliferation[48-49]. Although GBP4-focused functional assays have not been conducted to our knowledge, its most consistent prediction of higher PFS among GBP genes suggests that *GBP4* might be a promising gene for future experimentation.

### Limitations and Future work

Finally, the limitations of this study highlight opportunities for future work. Despite combining cohorts across two assays, the sample size of 140 remained below the recommended minimum of 200 for predictive modeling[2]. Feature filtering is also necessary when features greatly outnumber samples, but can remove clinically relevant genes. Therefore, expanding data sets by integrating cohorts across assays represents an important direction of future work that will likely substantially improve predictive modeling results. While matched pre-post data help mitigate patient- and assay-specific biases, such data are hard to obtain. Additionally, biomarkers/models ideally predict treatment response from pre-treatment expression alone. Because NACT is highly standardized in HGSC, models based on log2FC can still enable immediate action post-treatment. Our unique double-assay analysis also indicated that once limits of detection are addressed, assay-scaled expression values from NanoString and RNA-seq are directly comparable across samples. Therefore, building a model from combined pre-treatment RNA-seq and NanoString data using our preprocessing steps is feasible and could provide more immediate applicability. More broadly, the harmonization strategies, models, and resources available on our portal will be powerful tools for future data set incorporation and predictive modeling of HGSC and other cancers.

## Ethics Statement

Formalin fixed tissue blocks were acquired (COMIRB#18-0119) and a board-certified pathologist confirmed tumor tissue. This protocol was deemed exempt, as it used previously collected data, the information was not recorded in a manner that is identifiable, and the gained information did not change clinical decision making.

## Authors Contributions

HAT: Conceptualization, Data curation, Formal Analysis, Methodology, Software, Validation, Visualization, Writing - original draft, Writing - review & editing.

KRJ: Investigation, Resources, Writing - review & editing.

RJW: Methodology, Writing - review & editing.

LBVK: Writing - review & editing.

ND: Writing – review & editing

KB: Writing - review & editing.

MJS: Writing - review & editing.

RDD: Supervision, Methodology, Writing - review & editing.

AC: Conceptualization, Funding acquisition, Supervision, Methodology, Project administration, Writing - review & editing.

BGB: Conceptualization, Data curation, Funding acquisition, Methodology, Project administration, Supervision, Writing - review & editing.

## Acknowledgements

Thank you to Drs. Nicholas Adzibolosou and Ayesha Alvero from Wayne State University for their assistance in obtaining clinical metadata. We thank Dr. Luis Manso-Sanchez for his clarification on the meaning of their metadata. We thank the Biofrontiers Information Technology team for ensuring smooth computing and storage for our analyses.

