## Supplemental Material for "Cross-assay RNA modeling reveals cancer biomarkers"

April 29, 2026

### Supplemental Results

#### Harmonizing Counts between RNA-seq and Nanostring

##### Genes Considered in Harmonization

The NanoString platform uses probes considering a specific sequence subset of a gene's exons. This means that a NanoString count can apply to multiple isoforms of the gene. NanoString provides the RefSeq isoform to which a given probe is meant to capture so we first tried to link said isoforms to the most current RefSeq annotations. Of the 770 genes in the NanoString panel, 31 had exact NanoString naming matches to RefSeq, while 723 had isoform names matching except for the version number (e.g. A2M NM_000014.4 in Nanostring is NM_000014.6 in the current RefSeq).
The remaining 16 isoforms are not currently annotated in the most recent RefSeq version and therefore we instead linked the NanoString probe based on where it mapped in the genome (details in Methods). Three of these probes aligned to genes no longer existing in RefSeq (*CD45RA*, *CD45RB*, and *CD45RO*) but instead had to be manually linked to different isoforms of the same gene: *PTPRC*. Every one of the 770 probes mapped to the genome clearly only once, supporting NanoString's claim that NanoString is specific and accurate.
We attempted two methods for harmonizing counts: isoform-focused and exon-focused. NanoString annotation files provide the Refseq or ENSEMBL isoform of the gene used. Therefore, we first retrieved coordinates of gene isoforms matching to the names (Figure S5B). Probe sequences are about 100bp long and therefore best correspond to an exon rather than a complete isoform (i.e. collection of exons). Therefore, we also mapped probe sequences to individual exons for counting that might better match with NanoString counts. 765 probes successfully mapped to exons (details in Methods).

##### Normalization

Both NanoString and RNA-seq then rely on normalization to mitigate bias from a sample simply having higher general transcription levels, or sequencing/measuring biases of machines. NanoString uses relatively standardized normalization approaches, relying on positive and negative control probes along with pre-specified housekeeping genes to calculate normalization factors. Conversely, RNA-seq normalization approaches are widespread and often rely entirely on external factors (the length of regions being counted and the number of sequences/reads mapping to the regions being considered) rather than specifically calibrated internal controls. Due to the high variability of normalization approaches for RNA-seq and the fact that some data is only available with already normalized counts[1], we considered three normalization methods: Median-Ratio from DESeq2 (MR), Reads Per Kilobase of transcript per Million mapped reads (RPKM), and Transcripts per Million (TPM) (details in Methods, Supplemental Figure S5C.

#### Isoform-based Counts allow improved harmonization

Some biological explanation of isoform-counts being better regardless of limits of detection can be found by the fact that genes showing better harmonization with isoform-based counts tend to have longer probe-matching exons; therefore an exon is less comparable to the length of the probe (Supplemental Figure S10). We observed no clear feature differences for exon-improved counts nor between genes with improved harmonization from length-based vs non-length-based normalization (Supplemental Figure S11). Ultimately, isoform-based counts with length-based normalization, allowed more similar low-dimensional spaces of samples in NanoString and RNA-seq, regardless of removing genes below limits of detection (Supplemental Figure S12). Therefore, isoform-based counts seem to better harmonize with NanoString, likely by maintaining enough counts to overcome sampling and length-based biases. When considering highest-expressed isoforms vs NanoString-based isoforms, all cases with changed log2FC (outliers labeled in Figure 2E) were explained by expression levels changing between values already below our limits of detection and zero.Removing these cases led to $R^2$ values of 0.99 and below 1% of sample-gene combinations with different isoforms having changes in log2FC exceeding 1 and below 0.05% changing in sign. Of these, 1% problem cases, 60%-88% were explained by at least one of the transcription cases again being below the limit of detection.

#### Poor correlation for select samples in Matched Dataset

We wanted to make sure that poor or high correlation between NanoString and RNAseq of a gene was not being driven by a single outlier. Therefore, we recalculated spearman correlation coefficients after removing the top (maximum 2) samples with greatest distance from the line of best fit. Some genes significantly increased in correlation when only removing one sample (from spearman coefficients of 0.5 to greater than 0.85), regardless of harmonization approach. When looking into these genes, we saw that the outlier sample in each case was well within the range of expression (therefore not simply caused due to having expression levels not considered by other samples) and was almost always one of two samples: P_09 and P_18. We then checked if these sample outliers were explained by poor QC. Both samples, however, had uniquely mapped read numbers comparable to that of the other samples (31.6M for P_09 and 30.3M for P_18) and did not show any clear differences in mapping, sequencing, or trimming compared to the other samples. These discrepancies could be caused by multiple factors. First, mutations in the patient might allow specific genes to be poorly mapped to the probe. Additionally, it is possible that exact samples of the FFPE tissue blocks used for sequencing and NanoString were different enough to cause these discrepancies. Because we wanted to have strict consideration for whether a gene had poor or strong correlation, without bias from one specific sample, we removed these two samples for final coefficient calculations. For final results, we used the number of genes with correlation trends (good, poor, mid, mixture) across methodologies before filtering. After removing the samples, the number of genes with good correlation across all increased (345 to 390) and poor correlation decreased (116 to 78). We saw limited changes to distributions of spearman correlations (average increase of 0.04 and 0.002 increase in median spearman correlation for across and within patients, respectively). When determining characteristics of genes with improved correlation, we did not include the outliers to ensure we had clean comparisons. Importantly, we performed all RNAseq-vs-NanoString analyses with and without the samples, with no changes to trends. The genes with the largest impacts from outliers include: *BLM/IRF2/RELA*, and *G6PD/HLA-F* changed correlations of $0.5 - 0.6$ to $\geq 0.85$ when removing P_09 or P_18, respectively; *FUT4*, *UBA7*, *IFI35*, *CLEC4E* saw vast improvements when removing P_09 and P_18. Full comparisons of with and without the outlier samples can be found at

RNA-Nano-Matched/Final Comparison/Evaluate Harmonization.ipynb

#### Relevance of Harmonization in single-assay predictive modeling

We wanted to assess if our harmonization insights have improved upon the interpretation of the top 20 predictable genes based on several different modeling approaches (AUROCs < 0.6) for the same NanoString data[2]. When training a model using only these 20 genes, we achieved comparable performance to our original model (T/V AUROC=0.84, hold-out=0.78), but wit as almost entirely reliant on two genes: *CD274* and *MTOR* (Supplemental Figure S19B and C). *CD274* (protein PD-L1) was also among the top 25 predictive frequencies in both our RNA-seq and NanoString analyses. All genes with independent predictive power (measured by AUROC of Tr/V and hold-out above 0.55) showed expression levels above the limits of detection (*CD274*, *PDCD1*, *MTOR*, *IL11RA*, and *RAD51*), with 7/20 genes showing median expression levels below the limit in most experiments (Supplemental Figure S19A and D). Therefore, our harmonization approaches clearly allow improved insight both for feature filtering and model interpretation.

#### Evaluation of Predictive Models for PFS

##### Feature Relevance

To help clarify if some features captured redundant signals, we assessed how well each model performed when removing each individual gene, a leave one feature out strategy. For the combined (NanoString+RNA-seq) model, removal of most genes independently led to small (< 0.05) changes to AUROCs of the hold-out patients, with the most severe changes coming from removing *EGR1* (decrease > 0.1), *CTSS* (increase 0.1), *JAK2*, and *ENO1* (both decrease around 0.05) (Supplemental Figure S17 Top). When used as the only gene in the model, most of the model genes had AUROCs only slightly above 0.5, with *CD40* having the strongest predictive power (hold-out AUROC 0.81) and *JAK2*, *ENO1*, *RRM2*, and *IDO1* following as still having AUROCs clearly above 0.5 for both the full validation/training set and hold-out set (Supplemental Figure S17 Bottom). For the RNA-seq model, removal of *ESYT3* and *CLK2* showed no change in model performance, indicating clearly redundant signals from these and other genes in the model. However, removing either *CDKL2* or *GBP4* caused sharp declines (> 0.15) in AUROCs in the hold-out set, while *DDX11*, *HCN3*, *GPR173*, and *AMOTL2* showed only a slight decline (> 0.05) (Supplemental Figure S22). These findings further support that these six genes contain non-redundant information in the model. Conversely, removing *SHROOM1* caused a decrease in training/validation AUROC but an increase in the hold-out AUROC, suggesting that the gene's patterns might be more population-specific or the coefficient is prone to overfitting (Supplemental Figure S22). Indeed, we were unable to find clear literature support for the *SHROOM1*'s relevance in ovarian cancer.

##### Model confidence reflects biology

We then confirmed that the model confidence (logistic probability) matched that expected given our data. The combined model clearly favored one of the classifications (probabilities < 0.4 or > 0.6) for 100 of the 140 patients and low confidence classifications (near the decision boundary, i.e. ±0.1 of 0.5) were enriched in samples with PFS closer to the cutoff of 12 months (Supplemental Figure S15). As with the combined model, most cases (84%) were high confidence (predicted probabilities from the model were outside 0.4-0.6) and lower confidence predictions (probabilities of 0.5 ± 0.1) were enriched in patients with PFS values close to the cutoff (12 mths) (Supplemental Figure S14).

**Models perform better than random chance after feature filtering**

Since we have a feature filtering approach before modeling, we wanted to estimate 1) if we might be missing key genes with predictive power in the model, and 2) what a fair "random guess" baseline might be to compare for our model given we already limit features. To do this, we trained 1000 models on randomly selected sets of two genes and recorded the AUROCs on both the full training/validation set and hold-out set. For the NanoString+RNA-seq models, Median AUROCs from these random combinations were well below what our final model reached: 0.6 for training/validation and 0.5 for hold-out (Supplemental Figure S16A). There were a total of 17/1000 gene sets where the training/validation set had AUROC above 0.6 (max of 0.71) and the hold-out set above 0.8, with two of the cases including one of our final model genes: *CD40*. For the RNA-seq only model of PFS, we again found that the AUROCs values were significantly higher than AUROCs acheived across 1000 models using random sets of 2 genes (median AUC 0.45 Tr/V and 0.5 Hold out) (Supplemental Figure S16B). Only ten of the 1000 models had AUROCs above 0.8 (max 0.92) in one of the datasets (Tr/V or Hold-out) and above 0.7 (max 0.75) in the other, with *HCN3* being in one of the models. There was some stochasticity to this process, but the general results were observed regardless of seed used.

**Literature supporting model genes.**

Descriptions and citations of literature support and model genes can be found in Supplemental Table 3[3-33].

##### Guanylate-binding genes

While *GBP4* showed the most consistent and strongest trends and the highest AUROCs when considering different GBP genes in the full model, other guanylate-binding genes still show comparable if not greater predictive power individually (Supplemental Figure S23). *GBP3/4* showed consistent trends across all tested assays but *GBP3* was not included in the NanoString panel (Supplemental Figure S23C). Similarly, when considering the 4 HGSC non-longitudinal micorarray datasets, *GBP4/5* showed consistently significant results across 2 cohorts while *GBP1/2* showed in only one, and *GBP3/7* in none (Supplemental Table 4).

### 1 Supplemental Methods

Code for all methods can be found at https://github.com/Hope2925/NanoString-RNAseq-HGSOC/

#### 1.1 Nanostring Analysis

Nanostring counts were achieved by using nSolver Analysis Software version 4.0.70 on MacOS. Lanes were flagged when 0.5fM positive control ≤ 2 standard deviations above the mean from negative controls. Background thresholding was done according to the geometric mean of negative control counts. Positive control normalization was done using the geometric mean with lanes flagged if normalization factors were outside the 0.3-3 range. Reference/housekeeping normalization was done using the geometric mean with lanes flagged if normalization factors were outside the 0.1-10 range. No samples/lanes were flagged. RLF files from nCounter PanCancer IO 360 panel were used with platform and annotation information available on GEO using GEO accession GPL27956.

#### 1.2 RNA-seq Analysis

Reads were mapped to the hg38 genome where NCBI RefSeq annotations were used (hg38 release GCF 000001405.40-RS 2023 03) for all analyses except comparison with scRNA-seq in which case GENCODE v25 (RRID:SCR 014966) annotation was used to match what was used for the scRNA-seq. Bams were created by using the RNAseq-Flow nextflow pipeline at https://github.com/Dowell-Lab/RNAseq-Flow under commit 97c703b using the options –genome id 'hg38', -profile slurm, and –forwardStranded. Versions hisat2/2.1.0 (RRID:SCR 015530) fastqc/0.11.8 (RRID:SCR 014583), and bbmap/38.05 (RRID:SCR 016965) were used. Bbduk was used to trim fastqs which were then checked via fastqc. Hisat2 was used to map to hg38. Counting was done using featureCounts using subread v1.6.2 (RRID:SCR 009803). If using DESeq2 median-ratio normalization, RNA-seq counts were normalized using size factors from according to 191 housekeeping genes (full list found at 05_Get_Final_Counts.ipynb).

RNA-seq read counts were also normalized using Reads Per Kilobase per Million mapped reads (RPKM) and Transcripts Per Million (TPM). To avoid overlapping counts between transcripts of the same gene, size factors were calculated at the gene level.

For RPKM, gene-level counts were first scaled by the total number of reads mapped to genes in a given sample: $RPKM_{i,s} = \frac{C_{i,s}\ x\ 10^9}{N_s\ x\ L_i}$ , where $C_{i,s}$ is the read count for gene *i* in sample *s*, $L_i$ is the gene length in base pairs, and $N_s$ is the total number of mapped reads in sample *s*. RPKM was calculated for gene, transcript, and exon features separately. TPM was calculated using length-normalized counts. First, reads per kilobase (RPK) were computed as $RPK_{i,s} = \frac{C_{i,s}}{L_i/10^3}$. Next, these RPK values were normalized to sum to one million transcripts per sample as $TPM_{i,s} = \frac{RPK_{i,s}}{\sum_j RPK_{j,s}} x\ 10^6$, where the sum in the denominator includes all genes in the sample. TPM values were calculated separately for gene, transcript, and exon features. Annotated genes not captured by Nanostring probes were counted as all exons corresponding to the gene (no single isoform), or the isoform most highly expressed in the experiment. For the matched RNA-seq and NanoString dataset, RNA-seq counts were filtered so that genes not also considered in Nanostring were only kept if the median TPM value across all 24 samples was above 1 (leaving 17,976 genes).

#### 1.3 Harmonizing Counts

The code needed to replicate these analyses are found in NanoString-RNAseq-HGSOC/RNA-Nano-Matched. We first tried to link the NanoString probes to the isoform most similarly representing them by 1) using the isoform annotated by the NanoString company for the probe, or 2) considering the longest isoform that uses the exon(s) to which the probe maps (Isoform Counting). We also counted just over the exon(s) to which the probe mapped for a gene (Exon Counting), hypothesizing that this may allow for cleaner harmonization in the event of changing isoform usage (Supplemental Figure S5B). We next considered three different normalization approaches for RNA-seq: Median-Ratio from DESeq2 (MR),

Reads Per Kilobase of transcript per Million mapped reads (RPKM), and Transcripts per Million (TPM) (details in Methods, Supplemental Figure S5C). Further details are below.

##### 1.3.1 Probe Mapping

To account for probes extending across multiple exons, the probe sequences for the 770 Nanostring genes were mapped to hg38 using HISAT2. These sequences were also blasted (https://blast.ncbi.nlm.nih.gov/, RRID:SCR_004870) against the database Refseq genome RS_2023_03 using highly similar sequence algorithm (megablast).

##### 1.3.2 Exon matching

HISAT2 mappings of probe sequences were overlapped with hg38 exons (RefSeq (RRID:SCR_003496) hg38 release GCF 000001405.40-RS_2023_03). If a probe sequence was completely within an exon, those exon coordinates were used. Otherwise, all exons with the probe overlapping were considered. Probes that mapped to multiple genes (6 total probes) due to overlapping exons were linked to the coordinates of the exon(s) for the same gene as Nanostring. A subset of PTPRC exons simultaneously mapped to the probes for CD45RB, CD45RO, CD45RA, and PTPRC. Only the PTPRC probe was kept to avoid the same exons being considered across multiple probes, and because the PTPRC probe covered all the exons. A GTF file was produced to assign multiple exon coordinates to a single probe for counting. The full code and results for this can be found in subfolder Preprocessing.

##### 1.3.3 Matching Isoforms between RNA-seq and Nanostring

To identify isoforms best matching to NanoString probes, we first saw if the RefSeq isoforms NanoString offered as annotation of the probes could be linked to the most recent RefSeq hg38 annotations that we mapped RNA-seq reads to. CD45RA, CD45RB, CD45RO, and PTPRC in the Nanostring samples map to different exons of PTPRC, with no clear isoform matching up (CD45RA and CD45B and CD45RO no longer exist in the most recent annotation). For sake of completion, Nanostring probes for PTPRC, CD45RA, CD45RB, and CD45RO were assigned to isoforms that used the exon in which the probe mapped most exclusively: NM_080921.4, NM_002838.5, XM_047426398.1, and XM_006711472.5 of PTPRC, respectively. The full list of isoforms linked to probes based on HISAT and BLAST results can be found at 05_Get_Final_Counts.ipynb.
All probes, if successfully mapping to the genome, only mapped once. *LILRA3* has the Nanostring isoform of NM 001172654.2 which maps to an alternative reference assembly due to large differences to primary reference chromosome sequences (chr19 ALT_REF_LOCI_9), (both according to the isoform name and BLAST). Therefore the gene was not easily integrated into RNA-seq analysis and showed 0 counts across all RNA-seq datasets.

##### 1.3.4 Determining similarity between RNA-seq and NanoString

Spearman correlation coefficients were calculated to capture how NanoString expression compared to RNA-seq expression. We considered correlation of gene expression across patients (1 coefficient calculated for each gene) or within patients (1 coefficient calculated for each patient). These calculations were done for NanoString expression compared to each of the six approaches described in the RNA-seq Analysis section individually for RNA-seq based isoforms and NanoString-based isoforms (12 total comparisons). A gene was considered to have poor correlation if it had a spearman correlation coefficient across patients < 0.65 and high correlation if ≥ 0.8. Notes on two outlier samples can be found in Supplementary Results, but had little impact on final trends. A gene was considered to be better captured by Isoform- or Exon-based counts if the spearman correlation changed by at least 0.1 across at least two of the three normalization approaches. A gene was considered to be better captured by a normalization approach if there was a change of at least 0.1 across both Exon- and Isoform-based counts.

For evaluation of correlation and similarity of expression after considering low expression and range genes, we did the following. Genes that had ranges below the limits of detection were removed. Expression levels below the limit of detection were replaced with NA (therefore not included in analyses) and genes with NA values in 10 or more patients (out of 24) were removed. This led to about 635-700 genes kept when using strict cutoffs and 702-730 genes kept when using loose limits of detection.

RV coefficients were used to assess how similar the relationships were between samples in low-dimensional space without directly integrating RNA-seq with NanoString. We calculated them using FactoMineR::coeffRV and only considering genes with no samples having expression levels below the limits of detection. We then compared how samples mapped in low dimensional space (using PCA R function prcomp) when combined, with different approaches of scaling. In all cases, we used z-score scaling. We only changed scaling for the NanoString and RNA-seq samples, seperately, before performing gene-based scaling in the full (NanoString and RNA-seq) data-set as is the usual action for PCA. We either did not scale, scaled per-patient, or scaled per-gene separately before combination. The analysis for scaling comparisons are found in Evaluate_Harmonization_Post.ipynb.

We also investigated whether normalization approaches led to biases, but very few genes saw systematic improvements to correlation due to normalization: 9 genes better with length-based methods (RPKM and TPM) compared to MR, 5 genes better with MR than length-based methods, 5 genes better in RPKM compared to both MR and TPM.

##### 1.3.5 Estimating limits of detection for RNA-seq and NanoString

First, to bin genes generally according to low expression or range, we split genes according to 10 quantiles of expression and range (highest-lowest expression) for each of the six harmonization approaches. If considering all six harmonization approaches at once, each measurement would serve as a gene-harmonization approach pair rather than simply a gene. To estimate limits of detection, we first plotted NanoString expression and RNA-seq expression (separately for each of the six harmonization approaches), with each gene colored by the spearman correlation coefficient. We observed the worse correlation appeared close to low expression levels for both NanoString and RNAseq, mostly below where the largest mass of genes clearly began to cluster around a line-of-best fit. We first estimated loose cutoffs, where the NanoString (or RNA-seq) expression level reached the first point of the large cluster of genes. Strict cutoffs were determined by similar logic except where the first point of the large cluster of genes that also contain Spearman correlation coefficients clearly above 0.65. Without technical replicates, these limits were estimated largely by visual clustering and are not expected to be used as official limits of detection. The final estimated limits of detection for harmonization can be found in Supplemental Table 2. The code for this analysis is found in Evaluate_Harmonization.ipynb.

#### 1.4 Predictive Modeling of PFS and NanoString/RNAseq

The code and results for this section, and an outline of the number of samples in each prediction category and cohorts are found under RNA-PrePost/Modeling/.

##### 1.4.1 Training, Validation, and Hold-Out Sets

To ensure performance on the hold-out set was not biased from the patients held having either extreme PFS (therefore likely easier to separate) or PFS closer to 12 months (therefore likely harder to separate), we selected patients based on the following. For RNA-seq vs NanoString prediction, the patients with the third lowest, third highest, and median PFS were used from each mini-cohort (N=18). For PFS prediction, we selected the patients with the third lowest and third highest PFS values within each mini-cohort (N=12). This left 128 patients left for training/validation. When only considering one assay type (e.g. RNA-seq), to maintain a hold out N of 12 rather than 6, we added patients with the median PFS of each category (PFS > 12*mths* and ≤ 12*mths*) (except for the Manso mini-cohort which only had 4 patients with PFS > 12*mths*).

##### 1.4.2 Feature Selection

Due to our relatively small sample sizes, including the full feature list of a minimum shared 770 or all expressed 18,000+ genes would likely cause overfitting. Therefore, we performed an initial feature selection to confine the feature space considered by a model to a maximum of 25 genes, as is often done in high-dimensional datasets[34], [35]. First, we performed analyses with and without filtering of genes with median log2FCs ≤ 0.5 to optionally focus on genes with clearer changes in expression. When considering the full RefSeq annotated genes, we removed any genes where the median expression was below 0.4 RPKM (right below strict limit of detection) in both classifications: patients with PFS ≤ 12*mths* and > 12*mths*. This would ensure we are not removing genes that have large changes in expression (e.g. from no expression to high expression) but instead only genes with expression levels difficult to ascertain consistently. This led to gene numbers declining from 18,035 genes to 12,974, and keeping 652 NanoString genes.

For choosing the genes considered in model validation from the bootstrapping approach, the 10-40 genes with the highest predictive frequencies excluding that of a chosen threshold after 390 bootstrapped samples (cumulative proportions remained stable after around 200 samples). The threshold was chosen based on a clear visual clustering of genes that was consistently obvious as shown in these jupyter notebooks 02_NanoRNAseq_Btsp.ipynb and 03_RNAseqFull_BtspCons.ipynb. The exact numbers and genes used for consensus feature selection can be found at the jupyter notebooks 01_NanoRNAseq_Cons.ipynb and 03_RNAseqFull_BtspCons.ipynb. Scaling of features showed little to no difference in results. Therefore, we reported coefficients without scaling for more immediately interpretable results.

##### 1.4.3 Assessing Model performance

Since we performed some initial feature selection, we wanted to ensure that the random baseline from which to compare model performance was indeed still about 0.5. Therefore, we randomly selected two genes from the genes that had log2FC above 0.5 1000 different times and evaluated model performance

using each of these subsets. We recorded AUROCs on the full training/validation set and hold-out set to evaluate the AUROCs for each of these when using random subsets of genes.
We also calculated sensitivity and precision. We did this either using the whole dataset, or after splitting patients based on whether or not the probability from the model was greater than 0.1 distance from 0.5. Briefly, low-confidence patients had probabilities from the model of 0.4-0.6 (model is almost randomly guessing) while high-confidence patients had probabilities either below 0.4 or above 0.6.
We also assessed how much each feature of the final model was contributing to the model (e.g. if there were features capturing redundant signal). To do so, we reperformed training, and recalculated AUROC on the training/validation and hold-out sets when removing the single gene from the model. Similarly, we evaluated the predictive power of each feature individually by reperforming training and evaluation using only the single gene in the model.

#### 1.5 Non-longitudinal Microarray

Code and results for this analysis can be found at RNA-scRNA-Microarray/SC_Micro/03_Format_Microarray_KM_curves.ipynb.
Metadata was downloaded from Precog (https://precog.stanford.edu/).
Univariate Cox regression was performed on each of the four Microarray cohorts and gene-probes, using both a categorical variable (low vs high expression) or treating the expression as a numerical variable. For categorical Cox regression and Kaplan-Meier curves, we split the patients within each cohort based on the median expression. In both cases we used ‘coxph()’ for the regression and ‘survfit()’ for the Kaplain-Meier curves from survminer v0.5.1, survival v3.8-3, survMisc v0.5.6, and KMsurv v0.1-6. All model genes assessed for survival regression were considered by probes in at least two of the microarray datasets.

#### 1.6 scRNA-seq

Code and results for this analysis can be found at RNA-scRNA-Microarray/SC_Micro/. scRNA-seq data was pseudobulked (at the count level) for samples as well as sample/cell cluster levels based on the previous annotations from Zhang et al[36] using SingleCellExperiment v1.26.0 and muscat v1.18.0 in R. are 13 total samples originating from the same patient and tissue with both sequencing from scRNA-seq and bulk RNA-seq. Pseudobulked counts were then normalized using counts per million (CPM) or DESeq2 v1.44.0 (RRID:SCR 015687, MR) to ensure results were not driven by normalization approaches. Due to the high sparsity of scRNA-seq data, log2FC was calculated as log2((Post Norm+1)/(Pre Norm+1)). A total of 18,921 genes had non-zero counts in both scRNA-seq and bulk RNA-seq paired samples.

#### 1.7 Gene-Network Analyses

Code and results for this section can be found at RNA-scRNA-Microarray/Network/.

##### 1.7.1 hdWGCNA

Before running hdWGCNA v0.04.09, a Seurat object v5.3.0 and Seurat v5.4.0 (RRID:SCR_016341) for the 11 patients was normalized (NormalizeData()), the top 3000 variable features were found (FindVariableFeatures), and the expression was scaled (ScaleData()) with variables “nCount RNA” and “percent.mt” regressed out using option vars.to.regress. PCA was then run using RunPCA(seurat, npcs =

50). To maintain reasonable N, we did not include Endothelial cells (only 79 cells), combined all epithelial cancer cells (EOC) into one group (new N of 8,806), combined dendritic cells (pDC, DC-1, DC-2) into "Myeloid APC" (new N of 1,297), and combined NK cells (N=1,744) with ILC cells (N=288) into "Innate Lymphoid" or "NK ILC." Harmony v1.2.4 was then run using the first 30 PCs of PCA and group.by.vars of "sample" and "patient id". Metacells were then built within each treatment phase (Pre or Post NACT) and cell type (with the following changes above). Briefly, genes expressed in at least 4% of cells were considered and then MetacellsByGroups() was run with groupings being based on the cell types and patients with minimums of 50 cells needed. Several cell types lost patient representation due to not having enough cells, with the exact lost patients noted in the jupyter notebook below. Topological Matrices (TOMs) were calculated based on TestSoftPowers() using a signed network and ConstructNetwork() with default parameters. Briefly, the TOM similarity scores measure the relative interconnectedness of two nodes (genes) by comparing their shared neighbors and direct expression correlations across samples, thereby being more robust to noise than correlation alone[37]. Code for this analysis can be found at 04_hdWGCNA_Networks.ipynb.

##### 1.7.2 WGCNA

Variance stabilization transformation from DESeq2 v1.44.0 was used to get normalized counts for all available bulk raw RNA-seq data for pre or post-NACT conditions (N=80 or N=83). Code for this step can be found in 04_hdWGCNA_Networks.ipynb These data were then used to run pre/post NACT specific network analyses with WGCNA (v1.73) to create signed TOMs, with genes being split into two TOMs to avoid memory surges. This split is determined internally by WGCNA to keep similarly networked genes together. Code for this step can be found at 05_WGCNA_Networks.R
Two different approaches were used to identify *GBP4* gene networks specific to a certain timing (shared, Pre, Post). Both approaches led to the same findings described in results. First, genes that had TOM similarities above 0.1 in only one timing were considered timing-specific, and those above 0.1 in both were considered shared. Since Pre networks tended to have overall higher connectivities, we then classified genes to more specifically consider this potential bias based on both scores and differences in scores. Genes shared between Pre and Post NACT networks had TOM similarity scores above 0.11 in both Pre and Post, and differences < 0.02. Genes only found in Pre networks had score differences above 0.08 or the gene was not considered in the Post TOM despite having a connectivity with *GBP4* above 0.12 in Pre. Genes only found in Pre networks had score differences above 0.03 or the gene was not considered in the Pre TOM despite having a connectivity with *GBP4* above 0.11 in Post. Gene enrichment analysis was done with clusterProfiler::enrichGO where the universe of genes considered was the 23,557 genes reaching the required expression levels for the TOM in either pre- or post-NACT conditions. Code for this analysis can be found at 07_GBP4_Networks.ipynb

##### 1.8.3 Comparing

For each gene, the spearman correlation of TOM similarity scores were calculated between bulk and cell-type networks at Pre and Post conditions. Only unfiltered genes in both cases were considered. Code for this analysis can be found at 06_Network_Compare.R

### Supplemental Figures

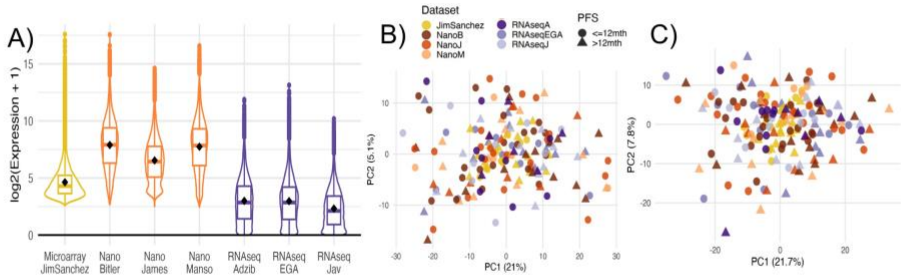


Figure S1: **Microarray shows a limited dynamic range with gene-scaling required for somewhat comparable levels.** A. Violin and box plots of normalized expression levels (log2 with pseudocount) of 731 genes shared across all assays and cohorts (RNA-seq normalization was RPKM). Black diamonds indicate means. B. PCA from pre-NACT expression alone of all cohorts/assays when scaling per gene before combining assays. C. PCA from Post-Pre NACT log2FC of all cohorts/assays when scaling per gene before combining assays.

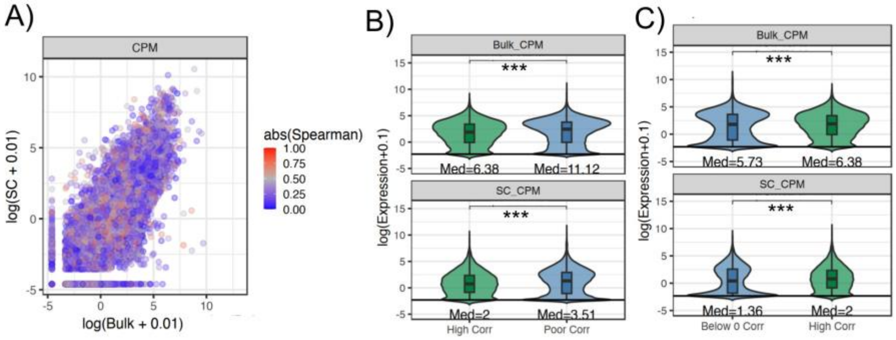


Figure S2: **Poor correlation between matched pseudobulked scRNA-seq and bulk RNA-seq is not explained by low expression genes.** A. Genes with low spearman correlation across matched pseudobulked scRNA-seq and bulk RNA-seq samples (N=13) (blue) are well spread across transcription levels predicted by both scRNA-seq and Bulk RNA-seq (x and y-axes). B. Genes with poor correlation have higher expression levels on average than those with high correlation. C. Genes with below 0 spearman correlation (N=) have lower transcription than those with high correlation (N). *** indicates p-value from Mann-Whitney test below 0.001.

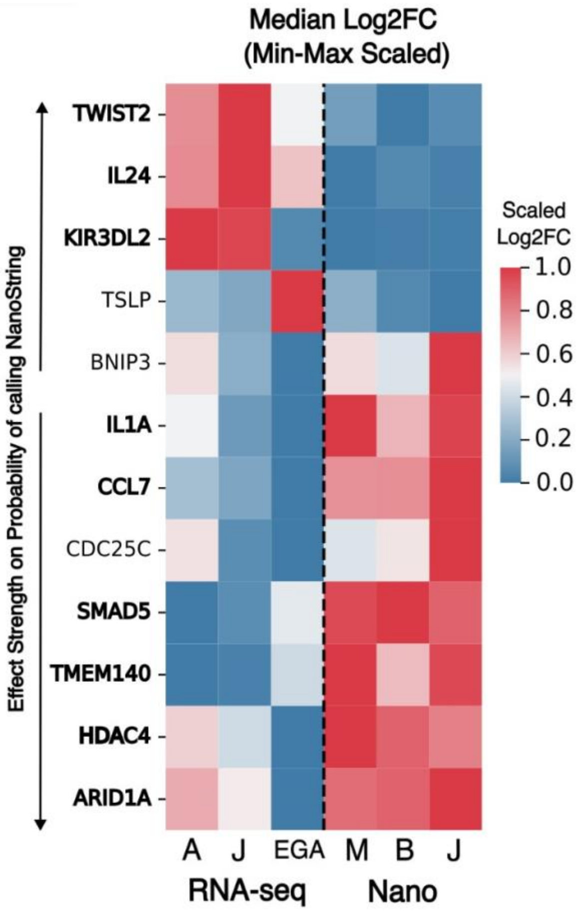


Figure S3: **Genes predictive of NanoString vs RNA-seq show consistent trends regardless of experiments.** Heatmap of final genes used as features in the model predicting NanoString from RNA-seq samples based on log2FC. Genes are ranked based on direction and effect strength (coefficient size) in the model calling NanoString. Bolded genes have the same19 trends across all experiments (NanoString vs RNAseq). Median Log2FC for each experiment are row-scaled with min-max scaling to more clearly visualize trends.

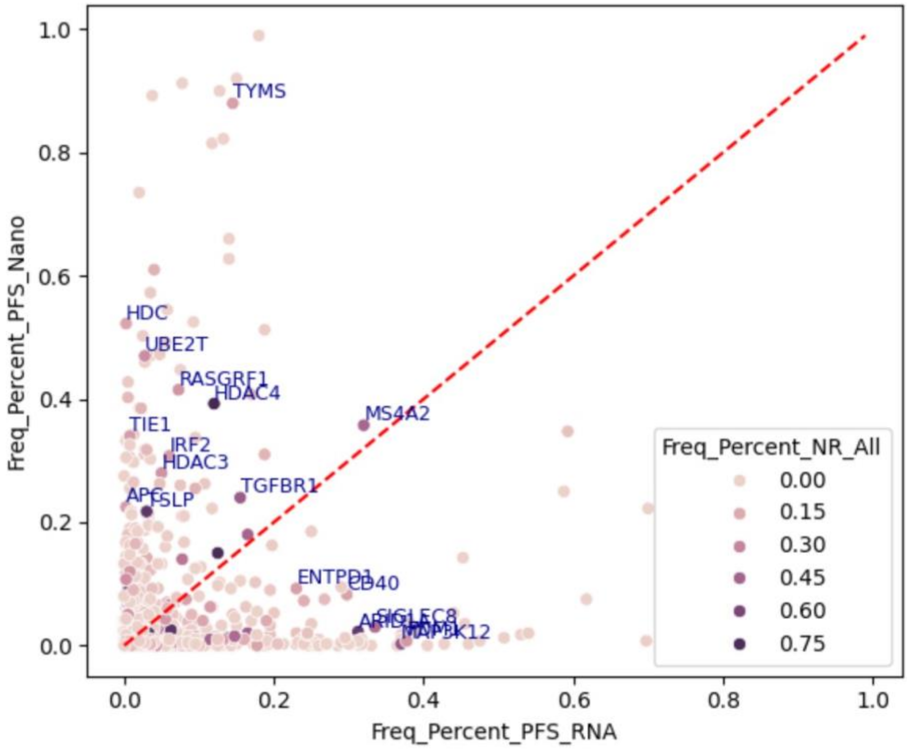


Figure S4: **Genes predictive of NanoString vs RNA-seq often shown high predictive power in either RNA-seq or NanoString.** Scatter plot of the cumulative frequency of a gene being within the top 100 predictive features across boostrapped samples (e.g. 1 means 100% of bootstrapped samples had the gene within the top 100 predictive features) for NanoString (y-axis) or RNA-seq (x-axis). Genes are then colored according to their cumulative frequency for bootstrapped samples in predicting NanoString vs RNA-seq samples (Freq_Percent_NR_All).

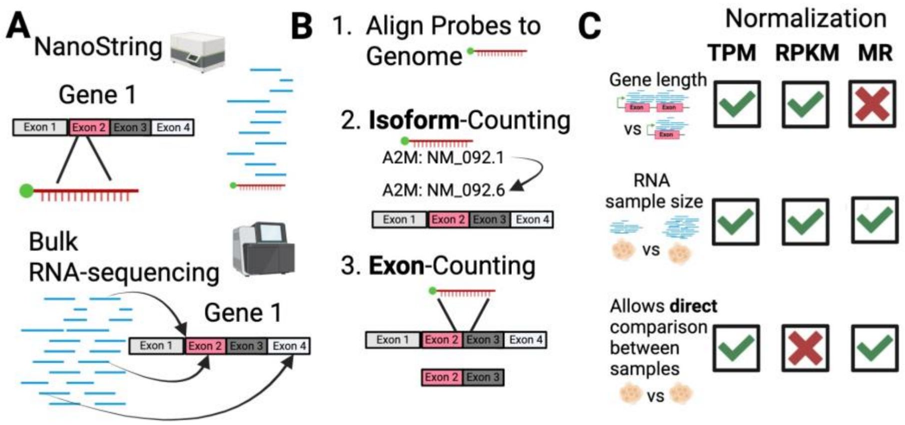


Figure S5: **Several preprocessing approaches were tested to harmonize counts across NanoString and RNA-seq A.** Visualization of what NanoString vs RNA-seq might capture based on a gene and its exons. NanoString will be confined to probe location while RNA-seq can map anywhere. **B.** We mapped probes to the genome and then used that to match a NanoString probe to either a given isoform (Isoform-Counting) or Exon(s) (Exon-Counting). **C.** Three RNA-seq normalization approaches were considered: Transcripts per kilobase million (TPM), Reads per kilobase million (RPKM), and DESeq2's median of ratios (MR).

Unlike TPM and RPKM, MR does not account for gene length. TPM and MR attempt to allow direct comparison between samples. All approaches consider library size. More details can be found in Methods.

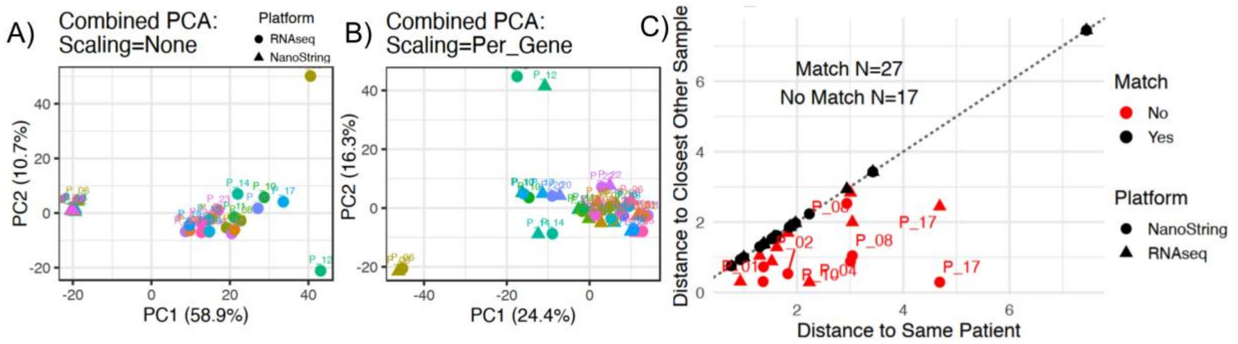


Figure S6: **RNA-seq and NanoString matched-samples are similar in low-dimensional space after gene-based scaling A.** PCA of RNAseq and NanoString matched samples without gene-based scaling. Patients are color coded and NanoString vs RNA-seq define most of the variance (PC1). NanoString and RNA-seq originally cleanly separate, with RNA-seq showing larger inner-variability. **B.** When performing gene-based scaling (z-score scaling for each gene across patients), patients are instead close together. **C.** When graphing the euclidean distance of a sample (NanoString or RNA-seq) to it's nearest other sample of the opposite assay (y-axis), most samples were closest to the sample of the same patient (27) (on the line so y=x).

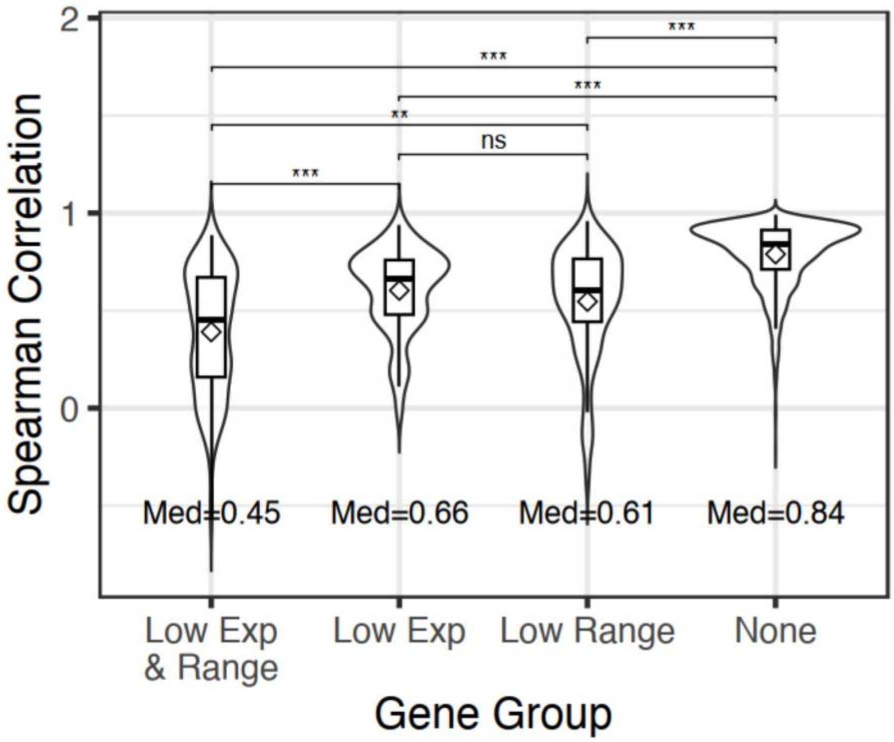


Figure S7: **Genes with low expression or low ranges of expression explain most of the poor correlation between NanoString and RNA-seq A.** Violin/Box plots of spearman correlation coefficients of genes between matched NanoString and RNA-seq split across four categories: those with both low range of expression and expression (Low Exp & Range), just low expression (Low Exp), just low range of expression (Low Range), or neither (None). Means are indicated by a diamond. Nonparametric Dunn's test was used for post-hoc pairwise comparisons and Bonferroni-adjusted p-values are indicated by ns > 0.05, ** < 0.01, *** <0.001.

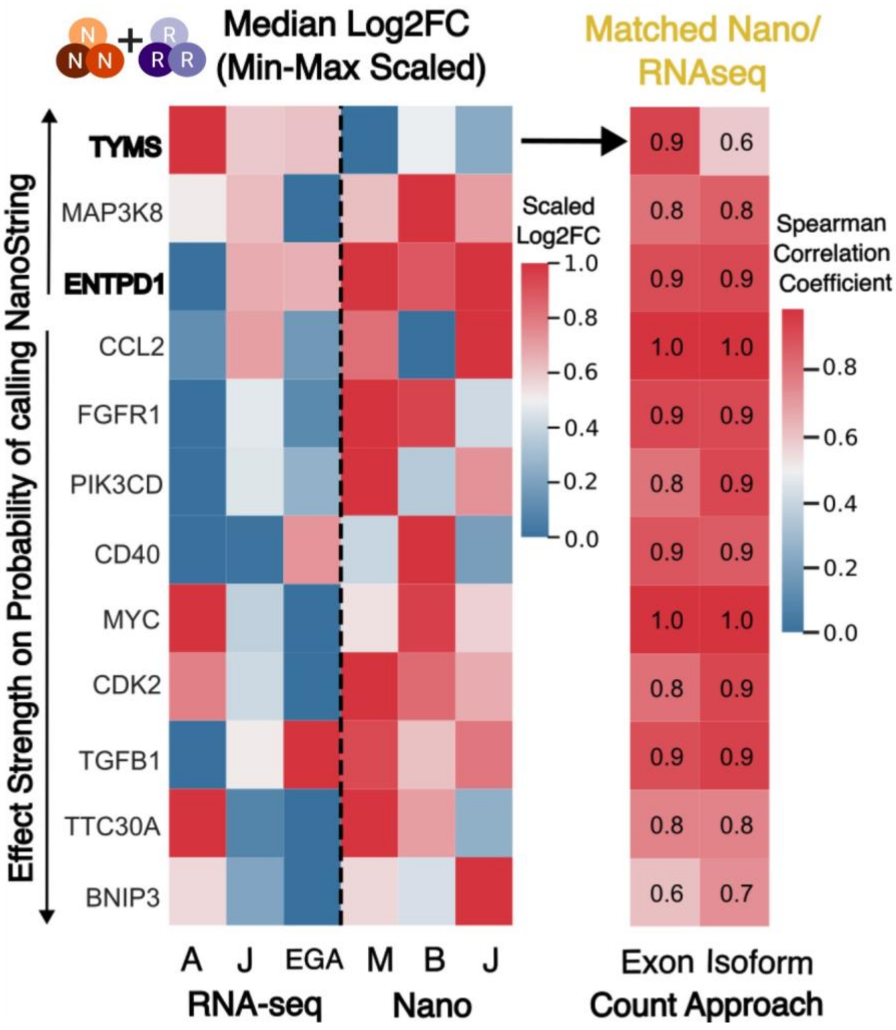


Figure S8: **Non low expression features marking differences between RNA-seq and NanoString log2FC are not consistent or improved by exon counts**. Heatmap of final genes used as features in the model predicting NanoString from RNA-seq samples based on log2FC. Genes are ranked based on direction and effect strength (coefficient size) in the model calling NanoString. Bolded genes have the same trends across all experiments (NanoString vs RNA-seq). Median Log2FC for each experiment are row-scaled with min-max scaling to more clearly visualize trends. Right shows heatmap of spearman correlation coefficients for the gene when comparing matched RNA-seq and NanoString used either Exon-based or Isoform-based counting.

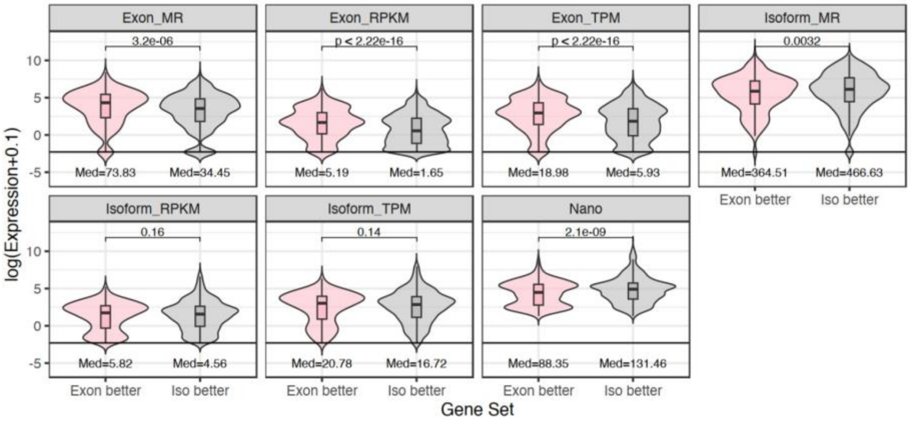


Figure S9: **Genes where exon counts have better harmonization have higher expression in exons than genes where isoform counts have better harmonization.** Expression levels (y-axis) for genes where exon-based counts show improved correlation between NanoString and RNA-seq than isoform counts (pink) compared to genes where isoform-based counts are better (grey). Exon-better genes have significantly higher expression in exons than isoform-better genes. There is no clear difference in expression levels for isoform-based counts. Pairwise comparisons were made using a Student's t-test with p-values shown. Med=median.

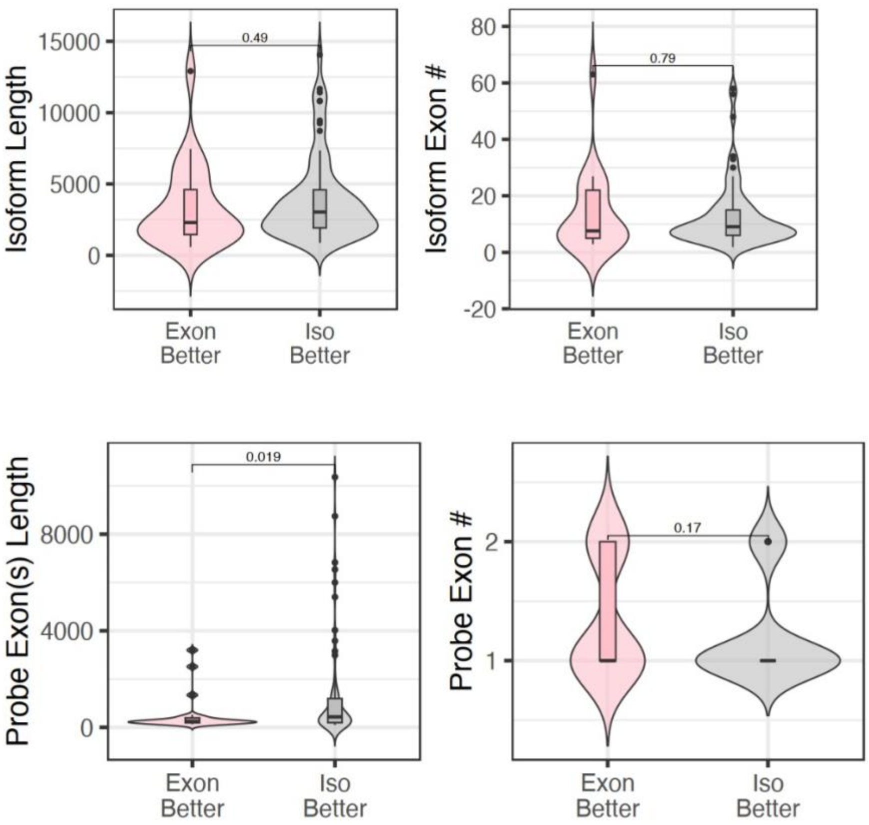


Figure S10: **Genes with better harmonization from isoform counts tend to have longer exons.** Genes with better harmonization between NanoString and RNA-seq from Exon-based counts (Exon Better - pink) or Isoform-based counts (Iso Better - grey) are compared for their isoform length, number of exons for the isoform (Isoform Exon #), lengths of the exons used for exon-based counting (Probe Exon(s) Length) and the number of exons to which the probe aligned (Probe Exon #). Pairwise comparisons were made using a Student's t-test with p-values shown.

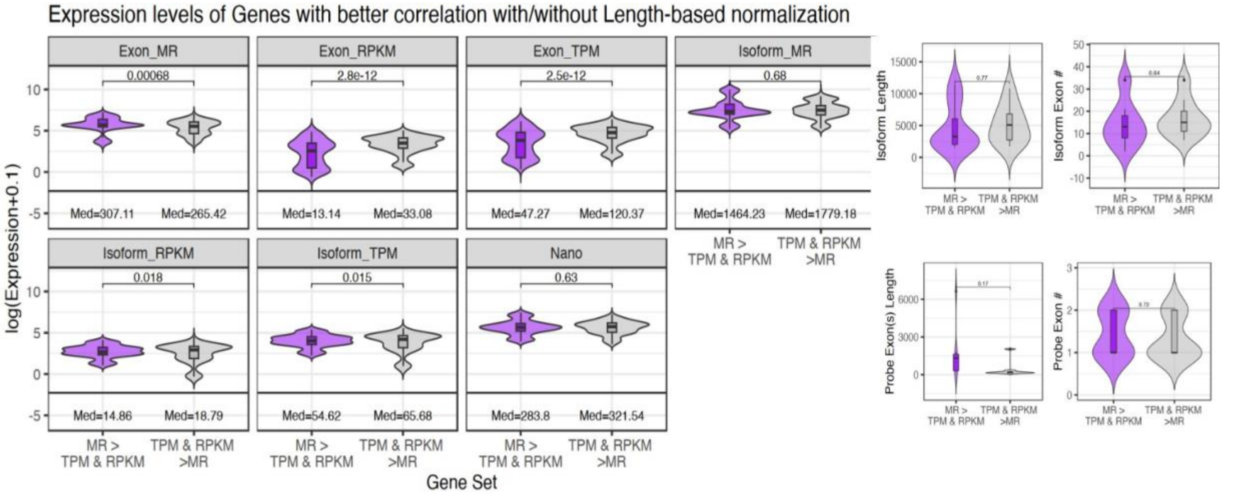


Figure S11: **There is no clear biological explanation for genes with higher harmonization via a normalization approach** Genes are split into whether non-length based normalization (MR) allowed better harmonization between RNA-seq and NanoString (purple) or if length-based normalization (TPM and RPKM) was better (grey). There is no consistent difference in Expression levels (left) or other features (right) (length of isoforms or exons, number of exons to the isoform or probes) between the two categories. Pairwise comparisons were made using a Student’s t-test with p-values shown.

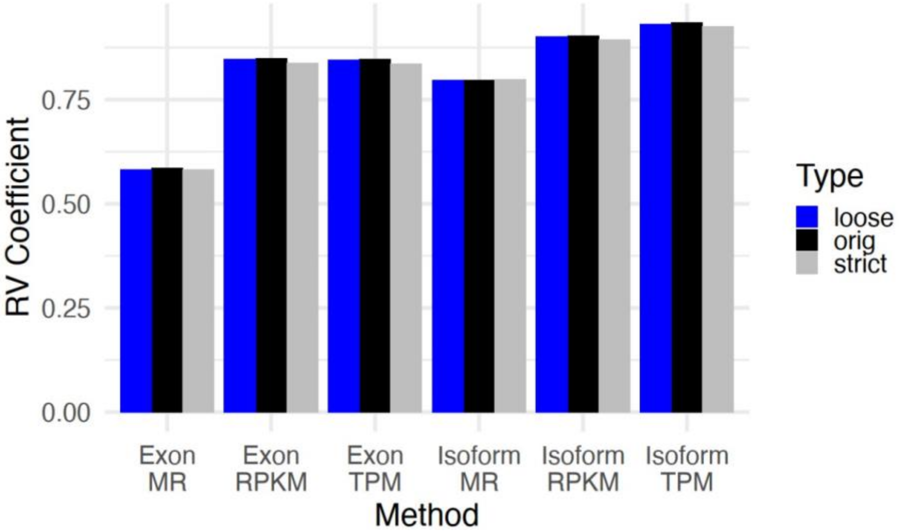


Figure S12: **Isoform and length-based normalization allow better harmonization in low-dimensional space between RNA-seq and NanoString.** RV coefficients (y-axis) measure the similarity of NanoString and RNA-seq in low-dimensional space when using the different harmonization methods (x-axis), where a higher coefficient indicates more similarity. Original refers to use of all 770 genes while loose and strict refer to when genes not passing the loose and strict limits of detection are removed.

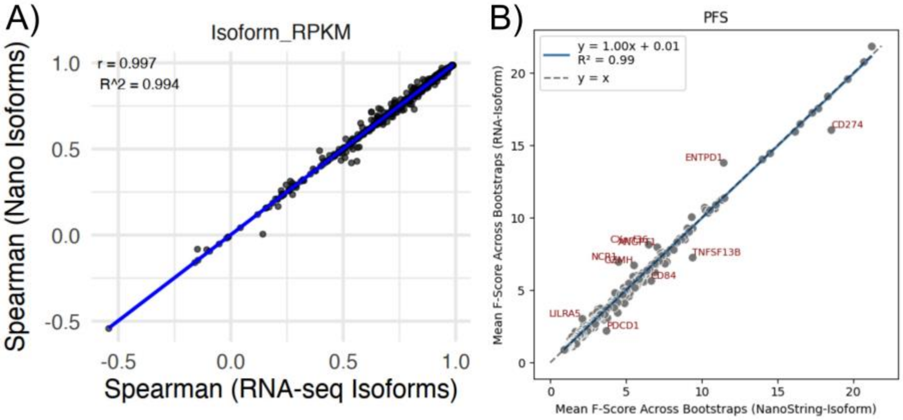


Figure S13: **Use of RNAseq-based isoforms has limited change on harmonization and predictive comparison between NanoString and RNA-seq A.** RNA-seq isoforms were chosen as the isoform most strongly expressed in TPM compared to other isoforms for a given gene. About 60% of isoforms changed from the original NanoString isoforms. Spearman correlations of the genes that changed isoforms between NanoString and RNA-seq are plotted when using NanoString-based isoforms (yaxis) and RNA-seq based isoforms (x-axis) cross ll three normalization approaches. The line of best fit along with r and $R^2$ values are indicated. **B.** Mean F-score of genes across bootstraps when using RNA-seq isoforms (y-axis) or NanoString isoforms (x-axis) are plotted with genes farther than 2 standard deviations from the fitted line highlighted.

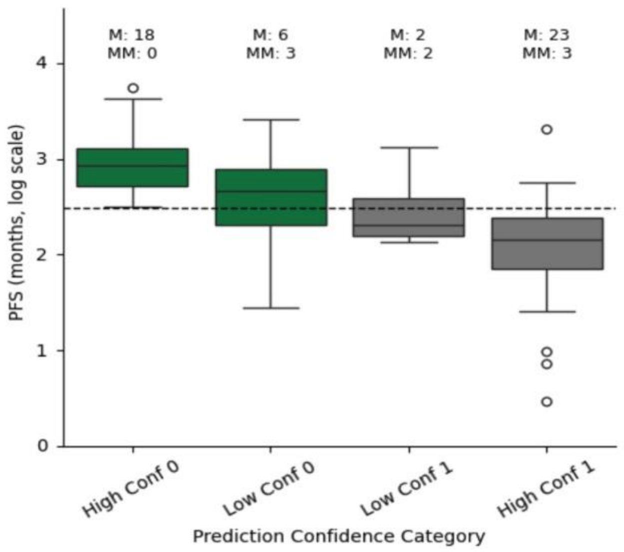


Figure S14: **Probabilities generated from the RNA-seq only model of PFS seem to reflect confidence of calls.** Patients were split into four categories based on whether the model called them PFS$\leq 12mths$ (1) with high confidence (probability above 0.6, High Conf 1) or not (Low Conf 1), or PFS$> 12mths$ (0) with high confidence (probability below 0.4, High Conf 0) or not (Low Conf 0). The true PFS are plotted on the y-axis with a line indicating the 12mth mark. Low confidence patients had PFS values closer to the cutoff of 12 and a higher ratio of mismatched call to reality (MM) to matches (M).

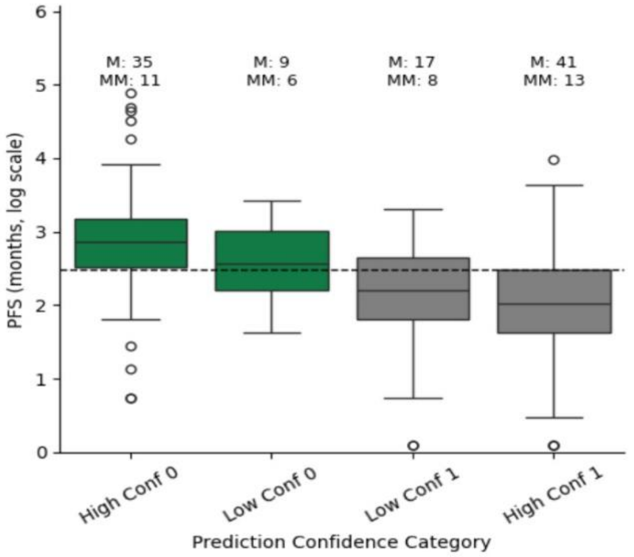


Figure S15: **Probabilities generated from the NanoString-RNAseq combined model of PFS seem to reflect confidence of calls.** Patients were split into four categories based on whether the model called them PFS$\leq 12mths$ (1) with high confidence (probability above 0.6, High Conf 1) or not (Low Conf 1), or PFS$> 12mths$ (0) with high confidence (probability below 0.4, High Conf 0) or not (Low Conf 0). The true PFS are plotted on the y-axis with a line indicating the 12mth mark. Low confidence patients had PFS values closer to the cutoff of 12 and a higher ratio of mismatched call to reality (MM) to matches (M).

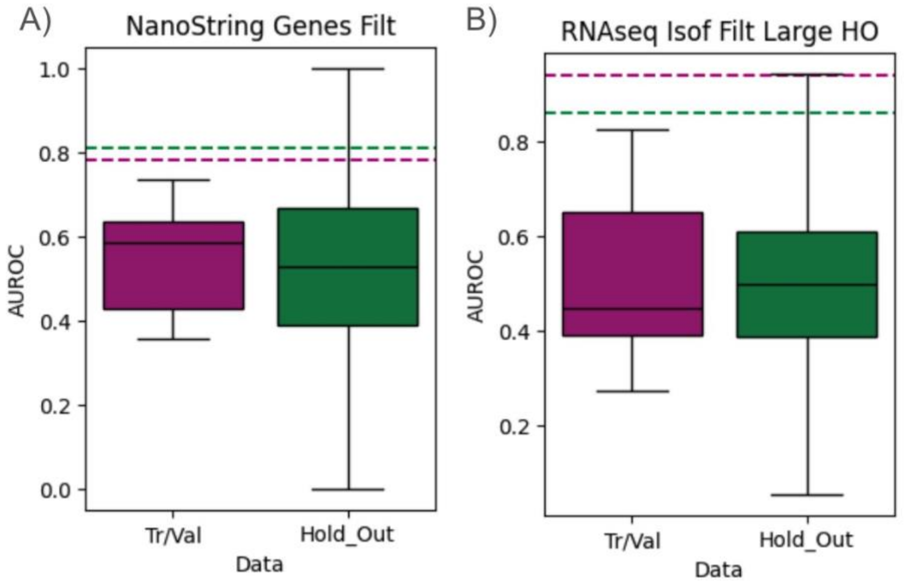


Figure S16: **Models using random sets of genes cannot recapitulate the same AUROCs of our final models.** Results from considering only the NanoString panel gene and combined dataset are in A while the full RNA-seq genes and RNA-seq dataset is in B. 1000 models were generated, each using random sets of 2 genes with median log2FC above 0.5 with AUROCs recorded for both the full training/validation set (All - purple) and hold out test set (Hold out - green). AUROCs generated from the final model from feature selection are shown as lines. For the combined datasets, all cases where the Hold out AUROC above 0.8 had All AUROC below 0.7 except IL2RA and CENPF (0.71 All and 0.82 AUROC). For the RNA-seq dataset, only 10 combinations had Hold out $AUROC > 0.8$ and All $AUROC > 0.7$ with the highest combination being 0.92 HO / 0.75 All when using genes GNB4 and CTSW.

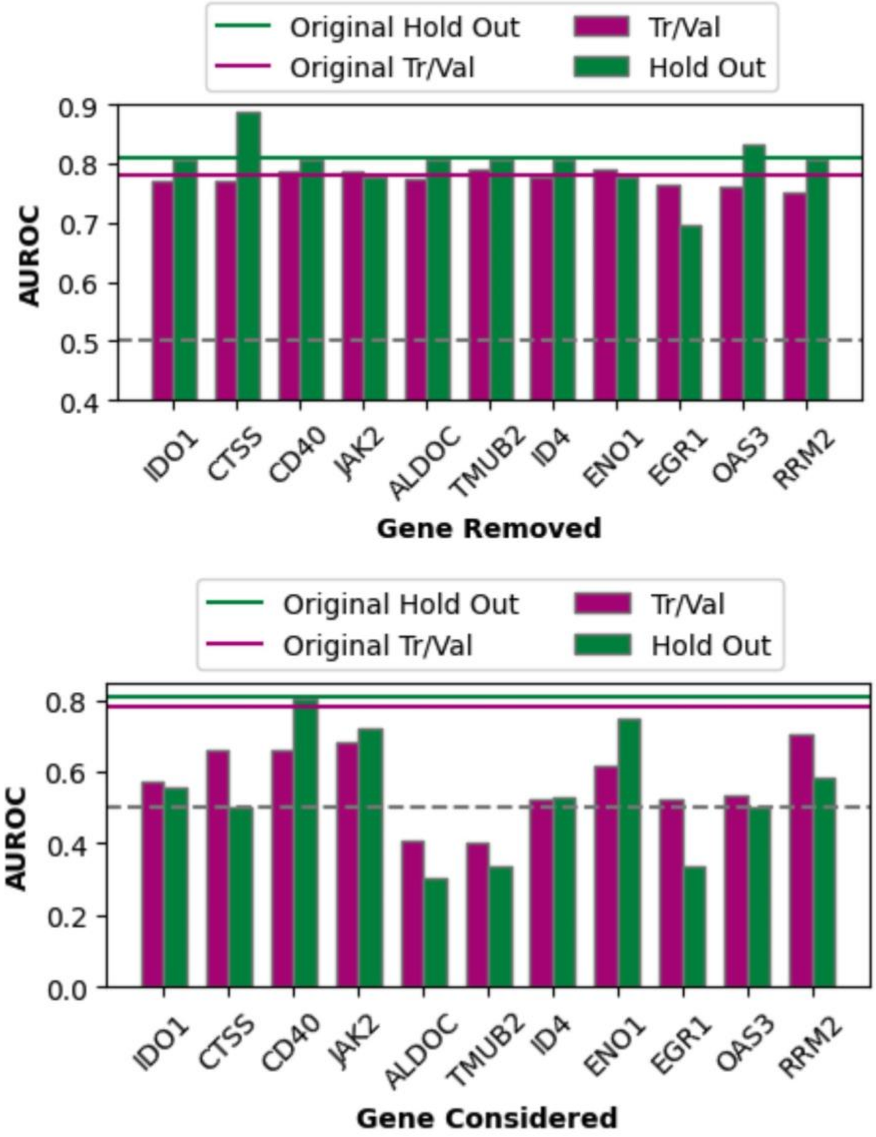


Figure S17: **There is some redundancy in features in the NanoString-RNAseq PFS model and varying individual predictive power across genes.** AUROCs of models including either all but one of the genes (top) or only a single gene (bottom) for the full training/validation set (All) or the Hold out set. A grey dotted line marks 0.5 AUROC (what is expected from random guessing) and solid lines indicate the AUROCs using the complete original model.

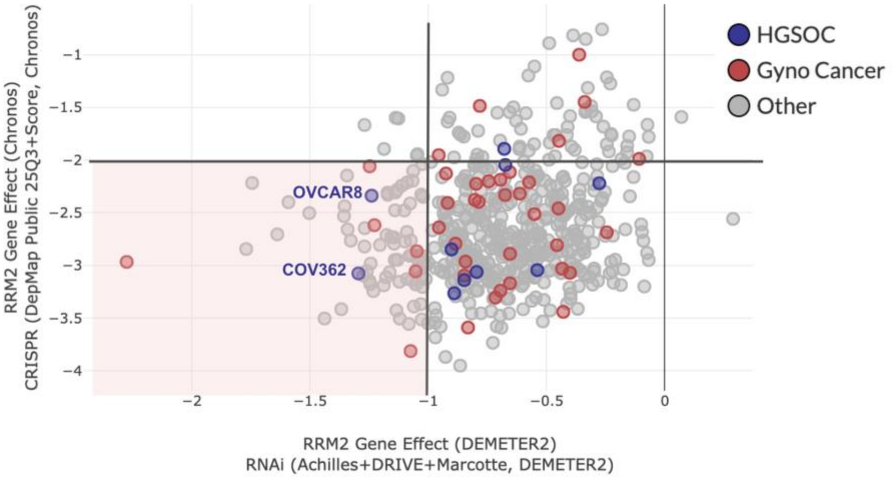


Figure S18: **RRM2 is a largely essential gene for ovarian cancer cell lines.** Scatterplot of gene effects of RRM2 from DepMap across cancer cell lines from CRISPR (y-axis) or RNA-interference (x-axis) where more negative values indicate more essential genes. DepMap suggested cutoffs of essentiality include -2 for CRISPR (Chronos algorithm[38]) and -1 for RNAi (DEMETER2 algorithm[39]) which are highlighted. Cancer cell lines from HGSC (purple) and gynecological cancers (red) are highlighted.

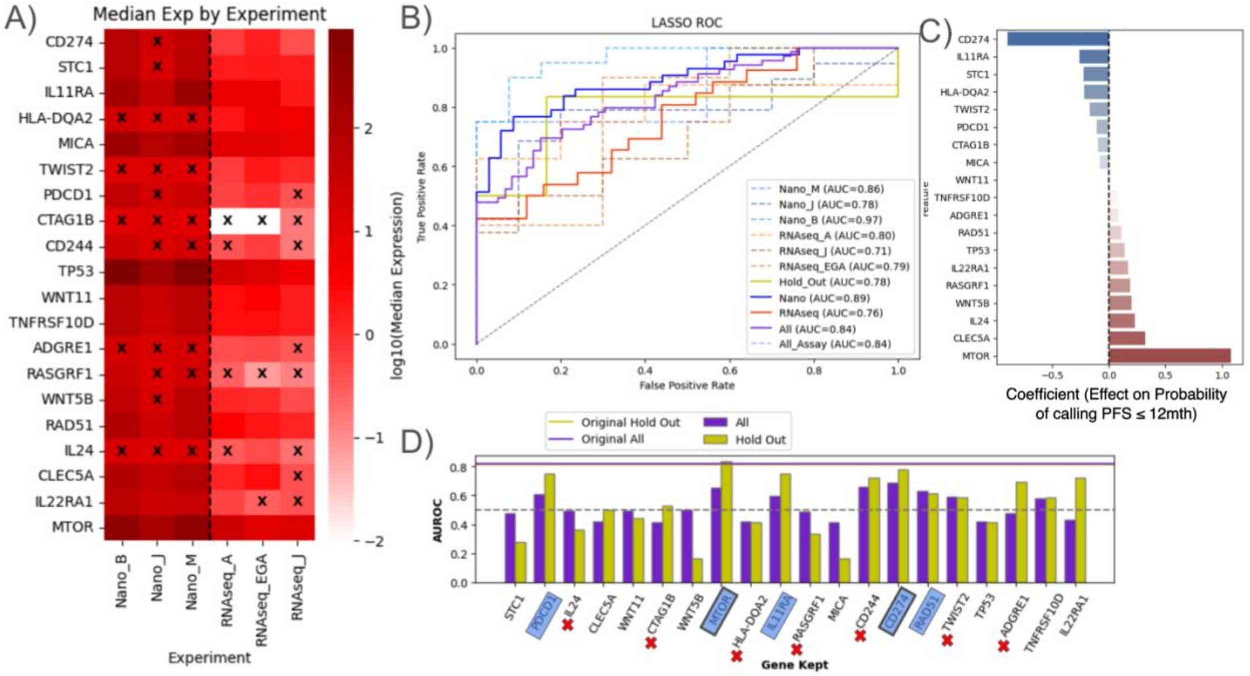


Figure S19: **Our harmonization approach informs about the top 20 ranked genes according to several different models and the same NanoString data** A) log10(Median expression) of genes across the minicohorts (experiments) where an X indicates the value is below the predicted limit of detection, B) AUROC plot for LASSO model using genes in C, C) Coefficient values for genes from the model, D) AUROC values for hold-out and training (All) set when each gene is considered alone in a model. Xs are placed by the genes with median expression levels below the limits of detection and blue highlights indicate where AUROCs of both training and hold out are above 0.6.

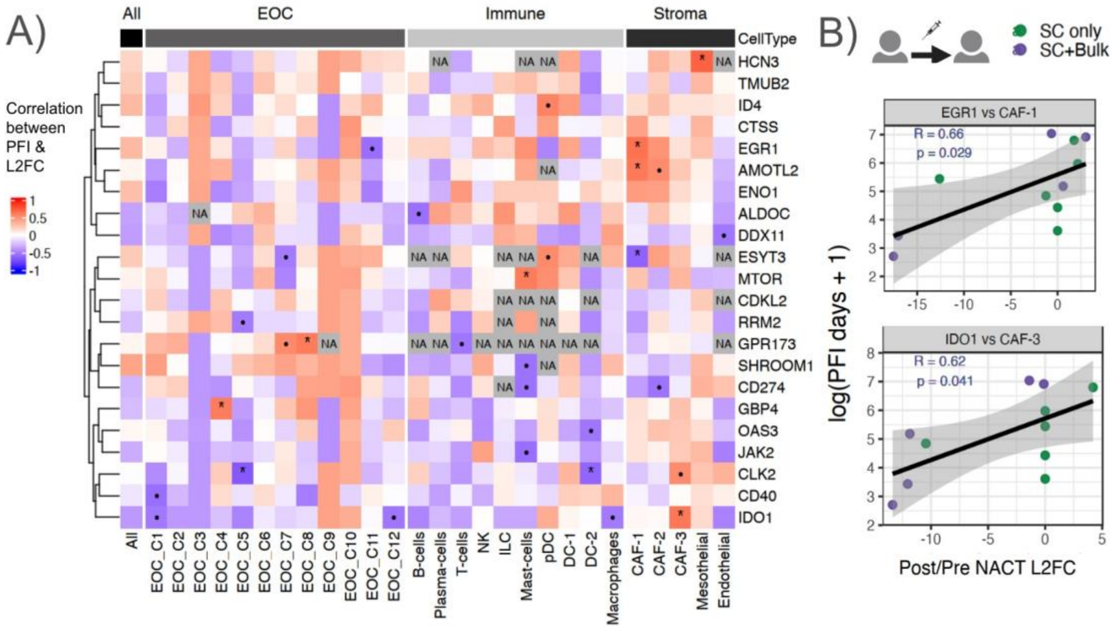


Figure S20: **Despite low correlation between pseudobulk scRNAseq and bulk RNA-seq, several model genes show significant correlation with PFI in a cell-state dependent manner A.** Heatmap of Pearson correlation coefficients between Post/Pre NACT log2FC of genes in the pseudobulked cell states and log(PFI+1) of 11 patients. An * indicates a Pearson correlation p-value below 0.05 and • below 0.1. *B.* Scatter plot of log2FC of gene in the noted cell type (x-axis) and log(PFI+1) of patients (y-axis) with the R-squared values noted according to best-fit line and dots colored by whether they have samples in Bulk (SC+Bulk) or just single-cell.

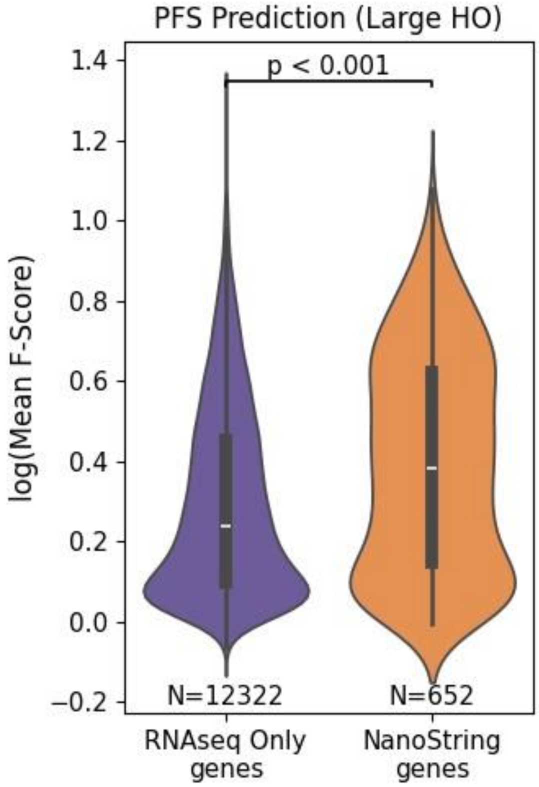


Figure S21: **NanoString genes had high predictive power compared to most RNA-seq genes.** Mean F-score of either genes only considered in RNA-seq (red) or in both NanoString panel and RNA-seq (blue) across bootstraps when using RNA-seq isoforms. Genes below count thresholds37 are not included. GBP4 and CD274 are NanoString panel genes that were consistently found in the top 25 genes with the highest predictive frequency.

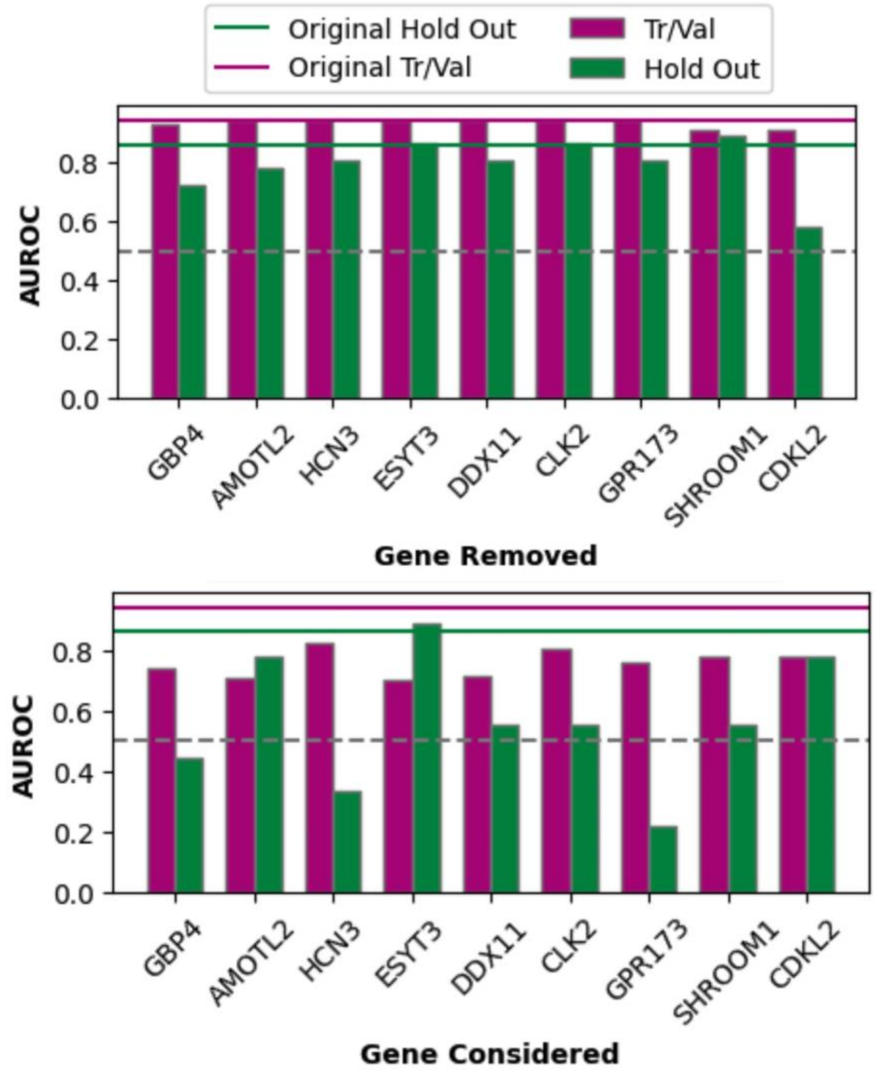


Figure S22: **There is some redudancy in features in the RNAseq PFS model and varying individual predictive power across genes.** AUROCs of models including either all but one of the genes (top) or only a single gene (bottom) for the full training/validation set (All) or the Hold out set. A grey dotted line marks 0.5 AUROC (what is expected from random guessing) and solid lines indicate the AUROCs using the complete original model. Removal of ESYT3, GPR173, or CLK2 had no impact on model performance despite having varying predictive power.

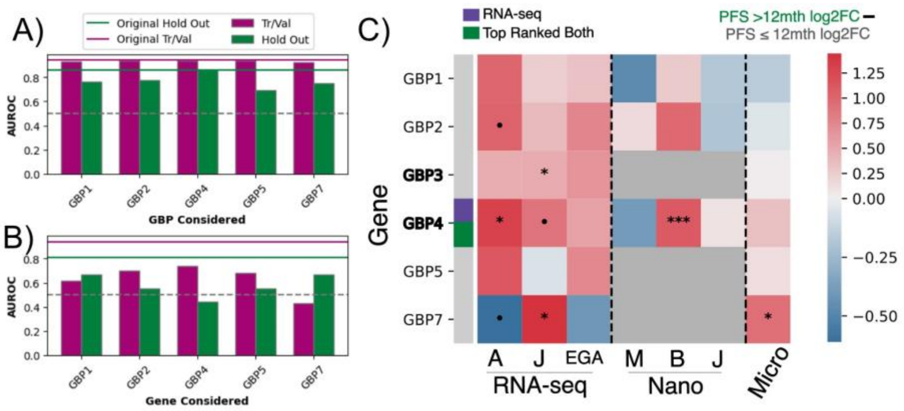


Figure S23: **Any one of the GBP genes might explain the GBP relevance A.** AUROC for the RNA-seq based model to predict PFS category on RNA-seq data when considering the different GBP genes in the dataset in the full model. **B.** AUROC when only using one of the GBP genes on RNA-seq data to predict PFS category. **C.** Heatmap comparable to Figure 4A of difference in median log2FC between patients with PFS ¿12mth and ≤ 12mth. • = p-value < 0.1, * < 0.05, *** < 0.001 for t-test (non-parameteric Mann-Whitney shows comparable results).

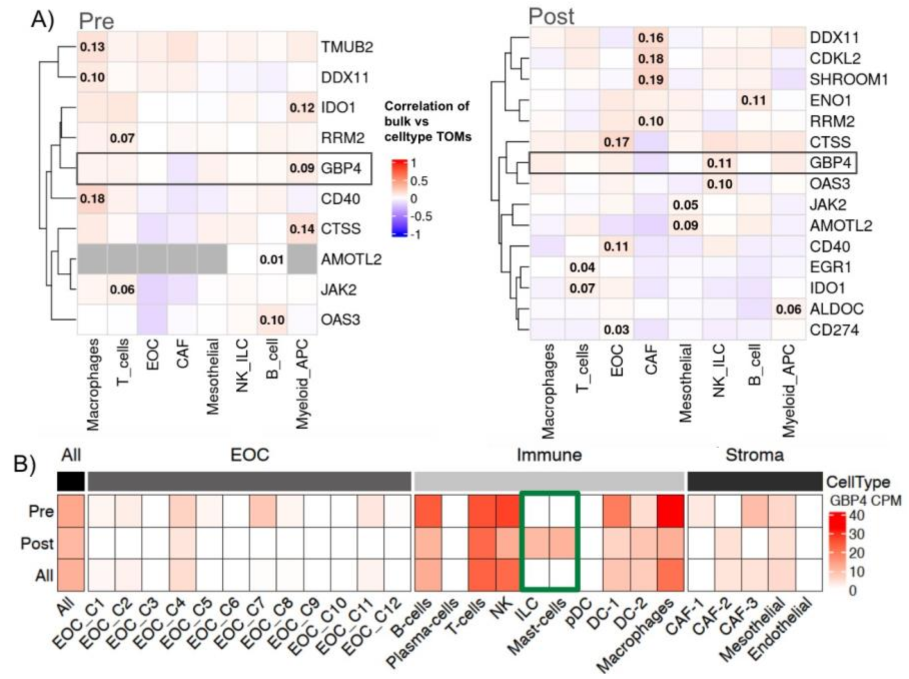


Figure S24: **Comparing GBP4 bulk expression within the context scRNA-seq A.** Spearman correlation of TOM similarity scores (edges) between model genes and other genes in Pre-NACT and Post-NACT conditions between bulk RNA-seq WGCNA and scRNA-seq hdWGCNA for cell-types. Grey or missing genes indicates that a gene did not pass filters for hdWGCNA for a given or any cell type, respsectively. Low correlation scores are expected given the consideration of 1000s of genes for each. **B.** Counts per million (CPM) within each pseudobulked cell-type from scRNA-seq Pre and Post NACT, or combined (All). Green boxes are around cell types that show an increase in GBP4 expression after NACT, as shows the highest correlation with improved PFS in the bulk model.
